# Suppression of tumor/host intrinsic CMTM6 drives anti-tumor cytotoxicity in a PD-L1 independent manner

**DOI:** 10.1101/2022.05.06.490847

**Authors:** Yiru Long, Runqiu Chen, Xiaolu Yu, Yongliang Tong, Xionghua Peng, Fanglin Li, Chao Hu, Jianhua Sun, Likun Gong

## Abstract

CKLF-like MARVEL transmembrane domain-containing 6 (CMTM6) has been identified as a regulator of membranal programmed death ligand 1 (PD-L1) stability and a factor associated with malignancy progression, but the effects and mechanisms of CMTM6 on tumor growth, as well as its potential for therapy, are still largely unknown. Here, we show that tumor CMTM6 increased with progression in both clinical patients and mice. Ablation of CMTM6 resulted in significant retardation of human and murine tumor growth dependent on T-lymphocyte immunity. Tumor CMTM6 suppression broke resistance to immune checkpoint inhibitors and remodeled the tumor immune microenvironment, as specific antitumor cytotoxicity was enhanced and contributed primarily to tumor inhibition. Further, without the PD-1/PD-L1 axis, CMTM6 suppression still significantly dampened tumor growth dependent on cytotoxic cells. Notably, we identified that CMTM6 was widely expressed on immune cells. T-cell CMTM6 increased with sustained immune activation and intratumoral immune exhaustion and affected the T-cell-intrinsic PD-L1 levels. Host CMTM6 knockout significantly restrained tumor growth dependent on CD8^+^ T-cells, and similarly, not entirely dependent on PD-L1. Thus, we developed and evaluated the antitumor efficacy of CMTM6-targeting adeno-associated virus (AAV), which effectively mobilized antitumor immunity and could be combined with various antitumor drugs. Our findings reveal that both tumor and host CMTM6 are deeply involved in tumor immunity with or without the PD-1/PD-L1 axis and that gene therapy targeting CMTM6 is a promising strategy for cancer immunotherapy.

**One Sentence Summary:** Even in the absence of the PD-1/PD-L1 axis, tumor or host CMTM6 deficiency can mediate cytotoxicity-dependent anti-tumor immune responses, allowing CMTM6 to be a novel target for scAAV-mediated oncoimmunology gene therapy and combination treatment.

## INTRODUCTION

Anti-programmed death ligand 1 (PD-L1) antibody treatment represents an inspiring clinical milestone for immune checkpoint blockade (ICB) therapies for malignancies (*1-5*). And the regulatory mechanisms of PD-L1 continue to be investigated in depth with the aim of solving the resistance and other problems (*6, 7*). In 2017, a novel PD-L1-interacting protein, CKLF-like MARVEL transmembrane domain-containing 6 (CMTM6), was identified as a critical regulator of PD-L1 membranal stability (*8, 9*). Colocalizing with membranal PD-L1, CMTM6 prolongs the half-life of PD-L1 on the cell membrane via obstructing the endosome-lysosome degradation pathway (*8*) and the ubiquitin/proteasome degradation pathway (*9*) of PD-L1.

*CMTM6*, located on chromosome 3, and seven other chemokine-like factor superfamily members were identified as new genes by homology search in 2003 (*10*). CMTM6 is an M-shaped four-transmembrane protein containing the MARVEL transmembrane domain (*10-12*). The PD-L1 regulation effect reported initially was subsequently confirmed in monocytes (*13*) as well as in other species of tumor cells (*14*). Extensive clinical analysis also found a significant positive correlation between CMTM6 and PD-L1 expression in tumors (*15-20*). Increased CMTM6 in tumors was associated with shorter overall survival (OS) and recurrence in patients with glioma, head and neck squamous cell carcinoma (HNSCC), melanoma, hepatocellular carcinoma (HCC) and triple negative breast cancer (TNBC), etc. (*18, 19, 21-24*). Remarkably, CMTM6/PD-L1 co-expression suggested a poorer prognosis (*20, 25-27*), but instead a better prognosis when ICB therapies intervened (*28-30*). However, some contradictory findings have also been reported, indicating a complex role for CMTM6 in clinical cancers such as non-small cell lung cancer (NSCLC) (*31-33*).

Research on the biological functions of CMTM6 is still in its infancy. CMTM6 promoted cisplatin resistance through acting with enolase-1 to activate Wnt signaling (*34*). The exosomal CMTM6 derived from oral squamous cell carcinoma cells augmented macrophage M2 differentiation by activating the ERK1/2 pathway (*35*). Studies also observed that CMTM6 inhibited p21 ubiquitination (*36*), genomic stability (*37*), and the mTOR pathway (*38*) to influence tumor cell growth, and acts on vimentin to affect tumor cell migration and invasion (*39*). And Hu-Antigen R (*40*) and neuropilin-1 (*41*) can affect the expression of CMTM6 at transcription and protein levels, respectively. In addition to the oncology study, researchers also found that Schwann cells optimize neural function by limiting axon diameter through CMTM6 (*42*).

In-depth and systematic researches are urgently required to elucidate CMTM6 functions in tumor progression. In this study, we extensively investigated the role and mechanism of human/mouse tumor CMTM6 as well as host cell CMTM6 in tumor immunity by CMTM6 ablation and explored the anti-tumor efficacy of gene therapy targeting CMTM6.

## RESULTS

### The association of CMTM6 expression with clinical malignancies

To systematically analyze the correlation between CMTM6 expression and clinical malignancy progression, we identified the CMTM6 expression differences between 33 cancers and normal tissues using the GEPIA tool (*43*) based on The Cancer Genome Atlas (TCGA) datasets and found that CMTM6 expression was significantly elevated in 10 cancers (Fig. 1A), such as colon adenocarcinoma (COAD) and pancreatic adenocarcinoma (PAAD). Not to be ignored, numerous other cancers also showed elevated CMTM6 expression trends. Then, the relationship between OS and CMTM6 expression was analyzed for 33 cancers, and high CMTM6 levels in three cancers, adrenocortical carcinoma (ACC), brain lower grade glioma (LGG) and PAAD, were found to significantly suggest poorer patient prognosis (fig. S1).

**Fig. 1.**
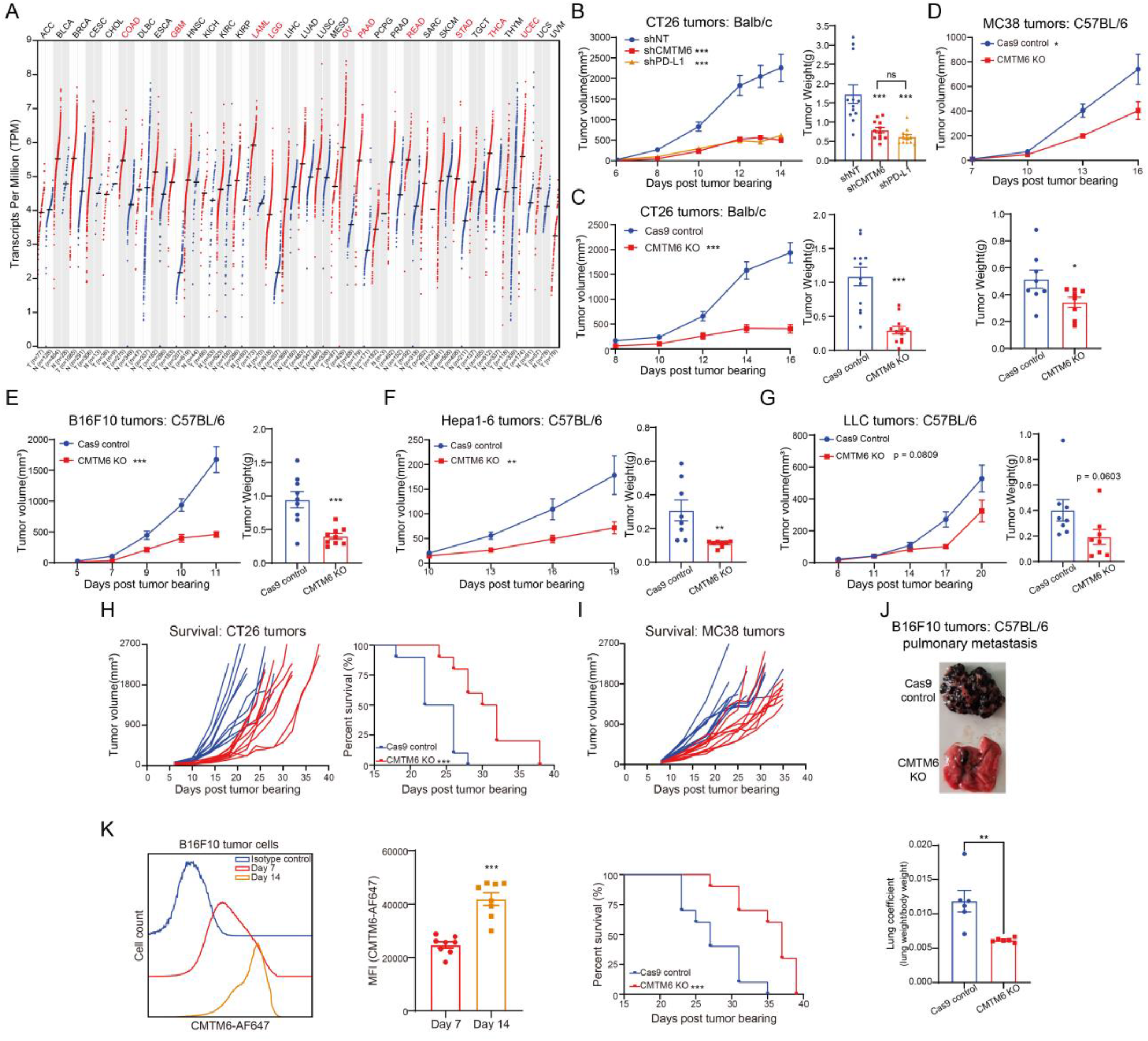
CMTM6 suppression attenuated tumor growth in syngeneic mice. (**A**) Human CMTM6 mRNA levels in different tumor types from the TCGA database were compared to normal tissues CMTM6 expression and were determined by GEPIA. The red font shows the tumors with a statistically significant difference in comparison by the Wilcoxon test. (**B**) Tumor growth kinetics of CMTM6-shRNA (shCMTM6) versus PD-L1-shRNA (shPD-L1) versus non-targeting shRNA(shNT) CT26 cells in Balb/c mice (n = 12). And tumor weights of mice in different groups are shown. (**C**) Tumor growth kinetics of CMTM6-sgRNA versus Cas9-control CT26 cells in Balb/c mice (n = 12). And tumor weight of mice in two groups are shown. (**D** to **G**) Tumor growth kinetics of CMTM6-sgRNA versus Cas9-control MC38 cells, B16F10 cells, Hepa1-6 cells and LLC cells in C57BL/6 mice (n = 8). And tumor weights of mice in different groups are shown. (**H**) Individual growth curves are shown for Cas9 control and CMTM6 KO CT26 tumors (n = 10). Right is the Kaplan-Meier plot of survival. (**I**) Individual growth curves are shown for Cas9 control and CMTM6 KO MC38 tumors (n = 10). Below is the Kaplan-Meier plot of survival. (**J**) A representative photograph of the whole lung is shown for pulmonary metastasis of Cas9 control and CMTM6 KO B16F10 tumors (n = 6). Below is the scatter plot of the lung coefficient (lung weight/body weight) for two groups. (**K**) Representative histograms show CMTM6 expression levels of B16F10 tumor cells measured by flow cytometry at different time points after tumor bearing. Right is the scatter plot of MFI (median fluorescence intensity) for CMTM6 in two groups (n = 8). The data are presented as the mean ± SEM. * p < 0.05; ** p < 0.01; *** p < 0.001; ns not significant by unpaired t test or one-way ANOVA followed by Tukey’s multiple comparisons test. The p values of survival plots were calculated by the log-rank test.

### CMTM6 deficiency affected murine tumor growth and metastasis

Next, to assess the effects of CMTM6 on tumor growth, we knocked out the *Cmtm6* gene in six murine cancer cell lines (CT26, B16F10, 4T1, LLC, Hepa1-6, and MC38), representing five tumor types, through the CRISPR–Cas9 system (fig. S2A). Using wound healing and proliferation assays, knocking out CMTM6 was found to not significantly alter the in vitro growth and migration rates of tumor cells (fig. S2, B to I). Consistent with earlier studies, CMTM6 deficiency significantly reduced cell surface PD-L1 expression in several mouse tumor cells with and without interferon-γ (IFN-γ) induction (fig. S3). For colorectal cancer (CRC) cell CT26, silencing CMTM6 showed a significant tumor suppressive effect in vivo with a 73.0% inhibition rate (IR) compared to the control group (Fig. 1B), which was comparable to PD-L1 knockdown (IR: 70.01%). Similarly, when CMTM6 knockout cells, including CRC cell CT26/MC38, melanoma cell B16F10, HCC cell Hepa1-6, and NSCLC cell LLC, were inoculated into syngeneic mouse hosts, their abilities to form tumors were also significantly attenuated in contrast to vector controls (Fig. 1, C to G). Moreover, the deficiency of tumor CMTM6 could effectively prolong the survival time of tumor-bearing mice (Fig. 1, H and I). In addition, tumor CMTM6 deficiency inhibited tumor metastasis in a B16F10 lung metastasis model (Fig. 1J).

### CMTM6 expression increased with tumor progression

With tumor progression, the dynamic changes of CMTM6 expression in vivo were monitored by flow cytometry (Fig. 1K). CMTM6 expression levels were 40.1% higher in 14-day melanoma cells than in 7-day melanoma cells. We then analyzed the clinical grading information available in the TCGA datasets for the aforementioned three cancers associated with patient prognosis, and found that all three cancers showed escalating CMTM6 expression with clinical disease development, particularly LGG and PAAD (fig. S4).

The above data illustrate that CMTM6 correlates with tumor progression and patient prognosis in clinical and mouse animal models. In pan-cancer, lowering CMTM6 in tumor cells effectively impedes tumor growth and metastasis in vivo.

### Correlation between CMTM6 expression and the immunophenotype of clinical tumors

To assess the impacts of CMTM6 on human tumor growth, we knocked out the *Cmtm6* gene in the human CRC cell line RKO by the CRISPR–Cas9 system (fig. S5A and Fig. 2A). In a humanized tumor model of RKO/human peripheral blood mononuclear cells (hPBMCs) mixed injection, ablation of CMTM6 significantly inhibited RKO tumor growth in vivo (Fig. 2A).

**Fig. 2.**
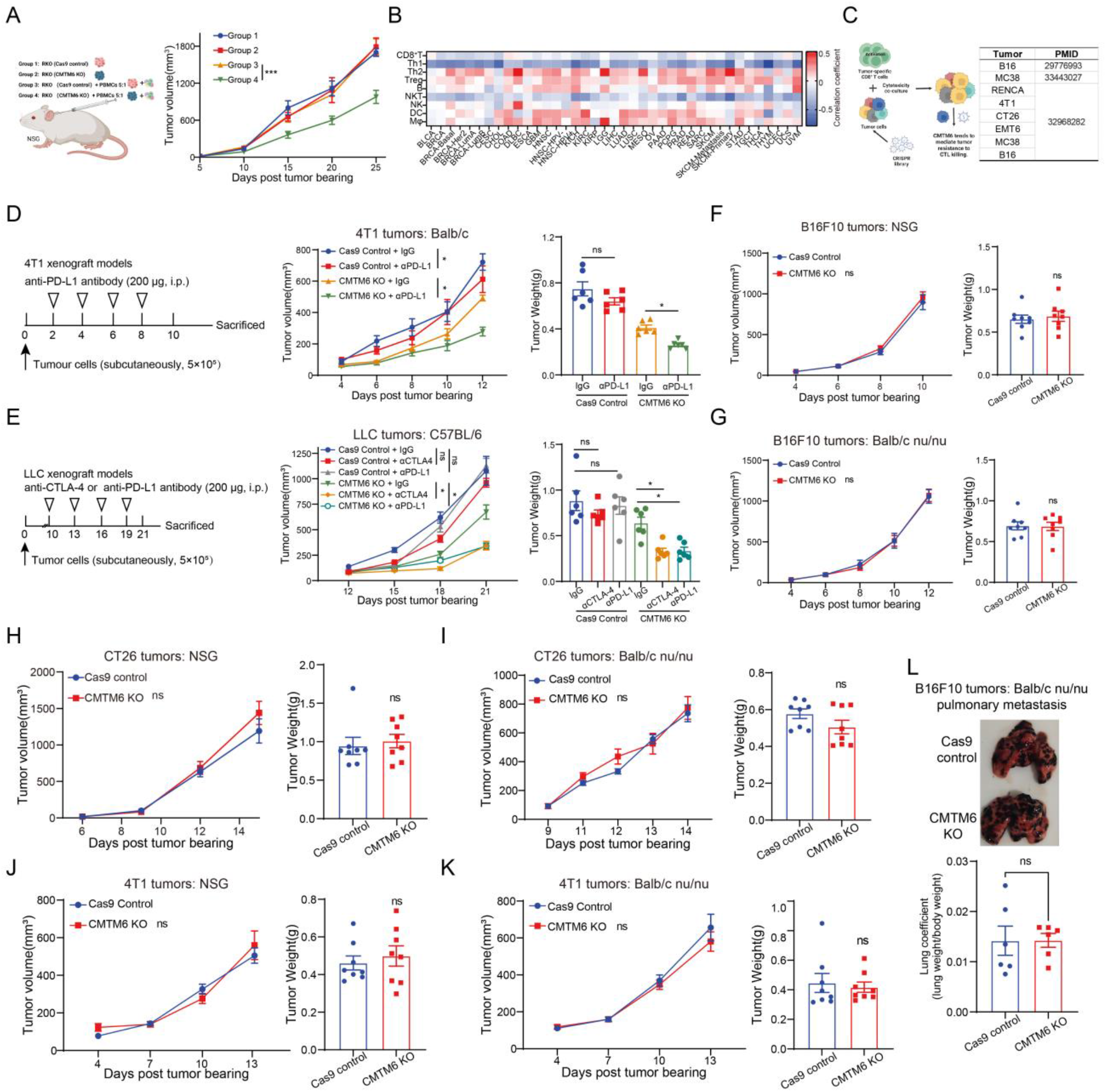
Suppression of CMTM6 overcame tumor resistance to ICB and attenuated tumor growth dependent on the host immunity. (**A**) Schematic diagram of grouping of Cas9 control and CMTM6 KO human RKO tumor cells with or without mixed injection of PBMCs. Tumor growth kinetics of four groups in NSG mice are shown (n = 8). (**B**) The heatmap shows the correlation of CMTM6 expression with immune cell infiltration level in different tumor types as indicated in the TCGA database and was determined by TIMER2.0. (**C**) Schematic diagram shows several CMTM6-deficient murine tumor cells promoting CTL killing from CRISPR library screening data reported in the literature. (**D**) Schematic of treatment schedules and dosing of anti-PD-L1 antibody against Cas9 control and CMTM6 KO 4T1 tumors in Balb/c mice (n = 6). Middle is the graph of tumor growth curves and the right is the scatter plot of tumor weight. (**E**) Schematic of treatment schedules and dosing of anti-PD-L1 antibody and anti-CTLA-4 antibody against Cas9 control and CMTM6 KO LLC tumors in C57BL/6 mice (n = 6). Middle is the graph of tumor growth curves and the right is the scatter plot of tumor weight. (**F, H, J**) Tumor growth kinetics of CMTM6-sgRNA versus Cas9-control B16F10, CT26 and 4T1 cells in NSG mice (n = 8). And tumor weights of mice in two groups are shown. (**G, I, K**) Tumor growth kinetics of CMTM6-sgRNA versus Cas9-control B16F10, CT26 and 4T1 cells in Balb/c nu/nu mice (n = 8). And tumor weights of mice in two groups are shown. (**L**) A representative photograph of the whole lung is shown for pulmonary metastasis of Cas9 control and CMTM6 KO B16F10 tumors in Balb/c nu/nu mice (n = 6). Below is the scatter plot of the lung coefficient for two groups. The data are presented as the mean ± SEM. * p < 0.05; *** p < 0.001; ns not significant by unpaired t test or one - way ANOVA followed by Tukey’s multiple comparisons test.

Interestingly, we found that this phenomenon was not observed in the highly immunodeficient mice NSG without hPBMCs injection, implying a connection between the effect of CMTM6 on oncogenesis and the immune system. Therefore, we then analyzed the correlation between CMTM6 expression and the immune cell infiltration in clinical TCGA tumors by the TIMER tool (*44*) and found that the expression of CMTM6 was positively correlated with the infiltration of T helper 2 (Th2) and regulatory T-cells (Tregs), and negatively correlated with the infiltration of CD8^+^ T-cells, Th1 and natural killer T-cells (NKT) (Fig. 2B). Moreover, in TCGA tumors with high CMTM6 expression, CMTM6 expression was significantly positively correlated with the expression of PD-L1, T-cell immunoglobulin mucin-3 (TIM-3), B7-H3 and other immune checkpoints (fig. S5, B to D). In addition, data from several genome-wide CRISPR library screenings (*45-47*) suggested that CMTM6 knockout in murine tumor cells promoted cytotoxic T-lymphocyte (CTL) killing to varying degrees (Fig. 2C). These data emphasize the relationship between host immunity, particularly T-cell immunity, and CMTM6 function.

### CMTM6 deficiency affected immune checkpoint inhibitor resistance

Subsequently, anti-PD-L1 antibody treatment-resistant 4T1 tumors and anti-PD-L1 antibody and anti-cytotoxic T-lymphocyte-associated protein 4 (CTLA-4) antibody treatment-resistant LLC tumors were selected to test whether CMTM6 knockout would affect the efficacy of ICB therapy. The results showed that CMTM6 suppression could restrain both tumors’ growth, and CMTM6-deficient tumors could respond to anti-PD-L1 antibody and anti-CTLA-4 antibody (Fig. 2, D and E). The anti-PD-L1 antibody inhibited CMTM6-deficient 4T1 with an IR of 43.4%. Anti-PD-L1 antibody and anti-CTLA-4 antibody inhibited CMTM6-deficient LLC with IR of 48.5% and 47.4%, respectively. These data suggested that tumor CMTM6 deficiency can reverse ICB resistance.

### Tumor inhibition by CMTM6 deficiency required host immunity

To further elucidate the involvement of the host immunity, we inoculated CMTM6-deficient and Cas9 control B16F10, CT26 and 4T1 tumor cells into NSG mice (Fig. 2, F, H and J). Results showed that CMTM6 deficiency had no effect on tumor growth in NSG mice. Furthermore, CMTM6 deficiency did not affect the ability of B16F10, CT26 and 4T1 tumor cells to form tumors in Balb/c nu/nu mice deficient in T cells (Fig. 2, G, I and K). In addition, the effect of CMTM6 deficiency to inhibit B16F10 tumor lung metastasis also disappeared in Balb/c nu/nu mice (Fig. 2L). These data suggest that host immunity, especially T-cell immunity, plays a decisive role in tumor suppression caused by CMTM6 deficiency.

### CMTM6 deficiency promoted antitumor cytotoxicity

Thus, we systematically investigated the involvement of host immunity in CMTM6 function. We first carried out a lymphocyte adoptive transfusion experiment (Fig. 3A). Tumor-infiltrating lymphocytes (TILs) from CMTM6 KO and Cas9 control CT26 tumors were isolated. CCK8 assays showed that TILs of CMTM6 KO tumors showed higher cell viability both at just isolation and after 24 hours of in vitro culture (fig. S6, A and B). The isolated TILs were then transfused back into the CT26 tumors. The results showed that compared to the PBS group, infusion of Cas9 control tumors’ TILs promoted tumor growth, while infusion of CMTM6 KO tumors’ TILs hampered tumor growth (Fig. 3B). And tumor volume and weight in the former group were more than twice as large as those in the latter group. On the other hand, the isolated TILs and CT26 cells were co-cultured in vitro. The results showed that TILs from CMTM6 KO tumors triggered apoptosis for 39.8% of CT26 and specific cytokine production, significantly higher than the Cas9 control tumors’ TILs (Fig. 3C and fig. S6, C to E). Notably, TILs from Cas9 control tumors were unable to produce IFN-γ when co-cultured with CT26, whereas TILs from CMTM6 tumors produced high levels of IFN-γ when co-cultured with CT26 (fig. S6D).

**Fig. 3.**
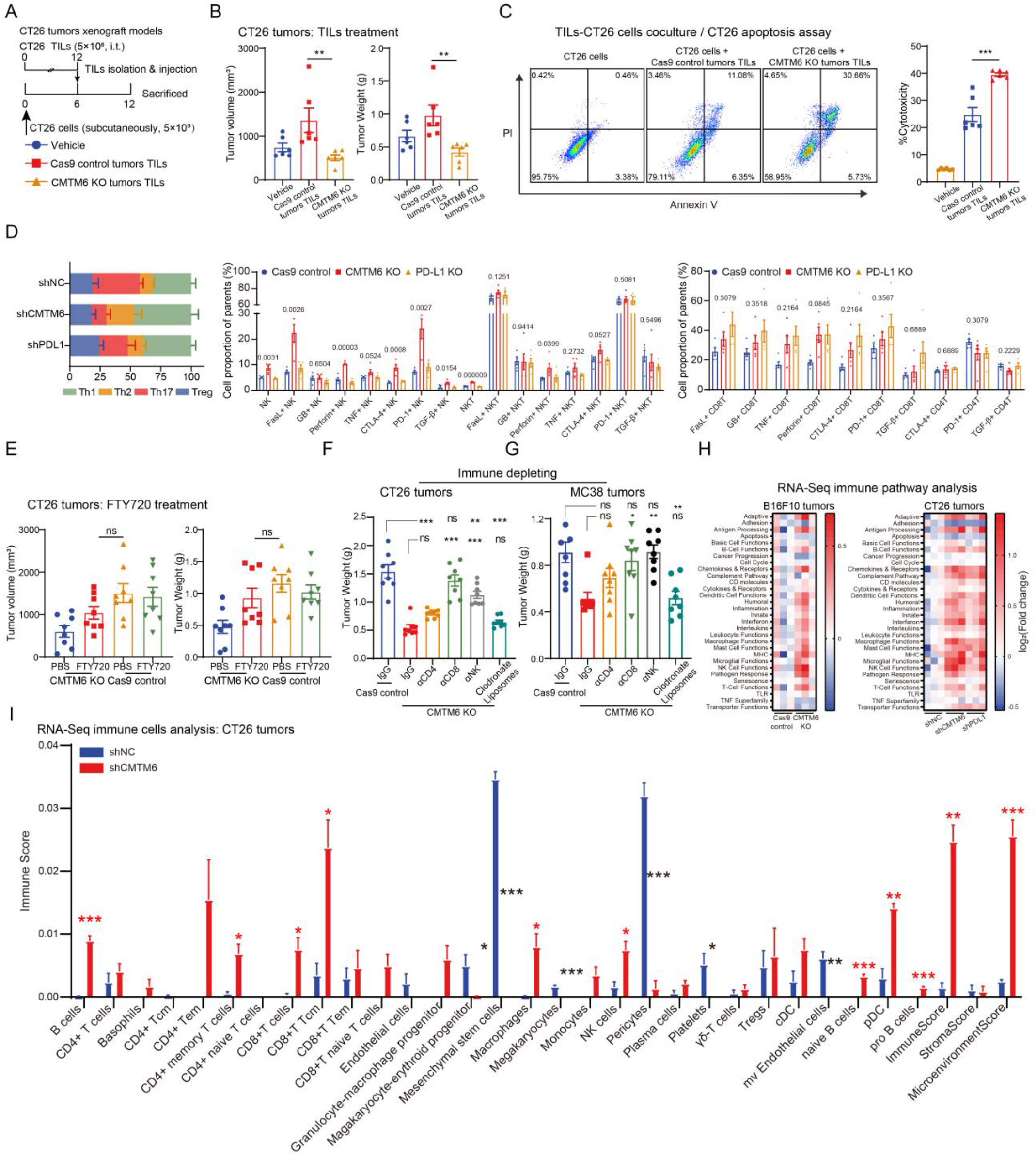
Suppression of CMTM6 enhanced intratumoral antitumor cytotoxic immune responses. (**A**) Schematic of TILs adoptive transfusion schedules. TILs isolated from Cas9 control and CMTM6 KO CT26 tumors were peritumorally injected into CT26 tumors. (**B**) The volume and weight of CT26 tumors in Balb/c mice injected with adoptive TILs are shown (n = 6). (**C**) CT26 cells and TILs were co-cultured at 5:1 ratio for 24 h. Representative density plots and scatter plot show the viability of CT26 cells in different coculture groups (n = 6). (**D**) At 12 days after tumor bearing, immune effector cells in shNC, shCMTM6 and shPD-L1 CT26 tumors were quantified by flow cytometry (n = 5). The percentages of specific CD4^+^ T, CD8^+^ T and NK cells are shown based on their respective markers. (**E**) Mice bearing Cas9 control and CMTM6 KO CT26 tumors dosed with FTY720. Tumor volume and weight are shown (n = 6). (**F**) Mice bearing Cas9 control and CMTM6 KO CT26 and MC38 tumors dosed with immune cell deletion reagents. Tumor volume is shown (n = 8). (**H**) Signature scores [defined as the mean log_2_(fold change) among all genes measured by RNA-Seq in the signature] for immune– associated processes are shown as heatmap (n = 3). (**I**) Immune cell infiltration analysis by xCell for RNA-Seq data (n = 3). The data are presented as the mean ± SEM. * p < 0.05; ** p < 0.01; *** p < 0.001; ns not significant by unpaired t test or one-way/two-way ANOVA followed by Tukey’s multiple comparisons test.

We next attempted to quantify immune effector cells in shNC, shCMTM6, and shPD-L1 CT26 tumors by flow cytometry (fig. S7 and fig. S8A). Immunophenotyping demonstrated that inhibition of tumor CMTM6 considerably promoted the effects of intratumoral cytotoxic cells, including NK cells, NKT cells, and CD8^+^ T-cells, significantly increased Th1 infiltration and reduced Th17 infiltration (Fig. 3D). The proportion of FasL^+^, granzyme B^+^, perforin^+^ and tumor necrosis factor-α (TNF-α)^+^ cytotoxic cells increased in CMTM6 KO tumors. Increased expression of some checkpoint molecules on effector cells may be due to higher activation levels of the immune system. Significant reductions in intratumoral PD-L1^+^ myeloid cell subsets and PD-L1^+^ tumor cells were observed (fig. S8B). Notably, CMTM6 suppression substantially improved NK cell function more than PD-L1 inhibition.

To determine the relative importance of different immune effector cells, we conducted a series of immune cell deletion experiments. The pivotal role of T-cell immunity in the involvement of CMTM6 in tumor immunity was reconfirmed by the inhibition of intratumoral infiltration of T-cells by FTY720 (Fig. 3E). We then found that CD8^+^ T-cells and NK cells played an absolutely dominant role in tumor suppression induced by CMTM6 inhibition in CT26 and MC38 tumors (Fig. 3, F and G). And deletion of CD8^+^ T-cells resulted in no significant difference in weight between CMTM6 KO and Cas9 control tumors in both models. The involvement of CD4^+^ T-cells and macrophages did not seem to matter, especially macrophages.

In addition, RNA sequencing (RNA-Seq) was applied to characterize the effect of tumor CMTM6 inhibition on the complex tumor environment (TME). The results showed that tumor CMTM6 inhibition affected multiple immune processes in both models, especially the adaptive immune process, interferon process, T-cell function, and NK cell function (Fig. 3H and fig. S9). In addition, xCell analysis of sequencing data showed that, consistent with the above results, infiltration of CD8^+^ T-cells and NK cells was significantly increased in shCMTM6 tumors (Fig. 3I). And platelets, pericytes, microvascular endothelial cells, and other tumor angiogenesis related cells decreased significantly.

Taken together, these results suggest that tumor CMTM6 inhibition remodels TME and mobilizes anti-tumor effect cells. Intratumoral cytotoxic cells and their specific tumor-killing effects mainly contribute to tumor suppression mediated by CMTM6 inhibition.

### CMTM6 deficiency inhibited tumor growth in absence of the PD-1/PD-L1 axis

Studies have established that CMTM6 can impact PD-L1 stability on tumor cell membranes, and our work also found that suppression of CMTM6 has similar effects on tumor immunity to PD-L1 inhibition. However, our results showed that CMTM6 suppression could only reduce membrane PD-L1 by less than half, but the tumor inhibition effect was comparable to that of PD-L1 inhibition, suggesting that CMTM6 may be involved in tumor immunity not only through modulation of PD-L1. Hence, we planned to investigate the biofunction of CMTM6 in PD-L1^-/-^ tumors.

Using wound healing and proliferation assays, knocking out CMTM6 was found to not significantly alter the in vitro growth and migration rates of PD-L1 KO tumor cells (fig. S10). When PD-L1/CMTM6 double-knockout CT26 cells were inoculated into syngeneic mouse hosts, their capacity to form tumors was significantly attenuated in contrast to PD-L1 KO and CMTM6 KO tumors (Fig. 4A). Moreover, 100% tumor regression was achieved in CT26 tumors of mice in the PD-L1/CMTM6 KO group. Two weeks later, the 10 survivors were rechallenged and monitored for tumor growth (Fig. 4B). Results showed that 7 of 10 CT26 tumors in the survivors achieved complete regression, and the weight of the 3 tumors left was significantly lower than that of the control group, suggesting that PD-L1/CMTM6 KO CT26 cells induced antitumor immune memory (Fig. 4C). Similarly, PD-L1/CMTM6 KO B16F10 tumors were significantly limited compared to PD-L1 KO B16F10 tumors (Fig. 4D). Furthermore, the MC38 model that was found to be unaffected by PD-L1 KO was selected. We found that only deletion of tumor CMTM6 inhibited tumor growth in WT C57BL/6 mice, while PD-L1 deletion did not play a role (Fig. 4E). Moreover, in PD-1^-/-^ mice, CMTM6 KO MC38 tumors were still limited in growth, and the tumor inhibition rate (40.9%) was the same as that of WT mice.

**Fig. 4.**
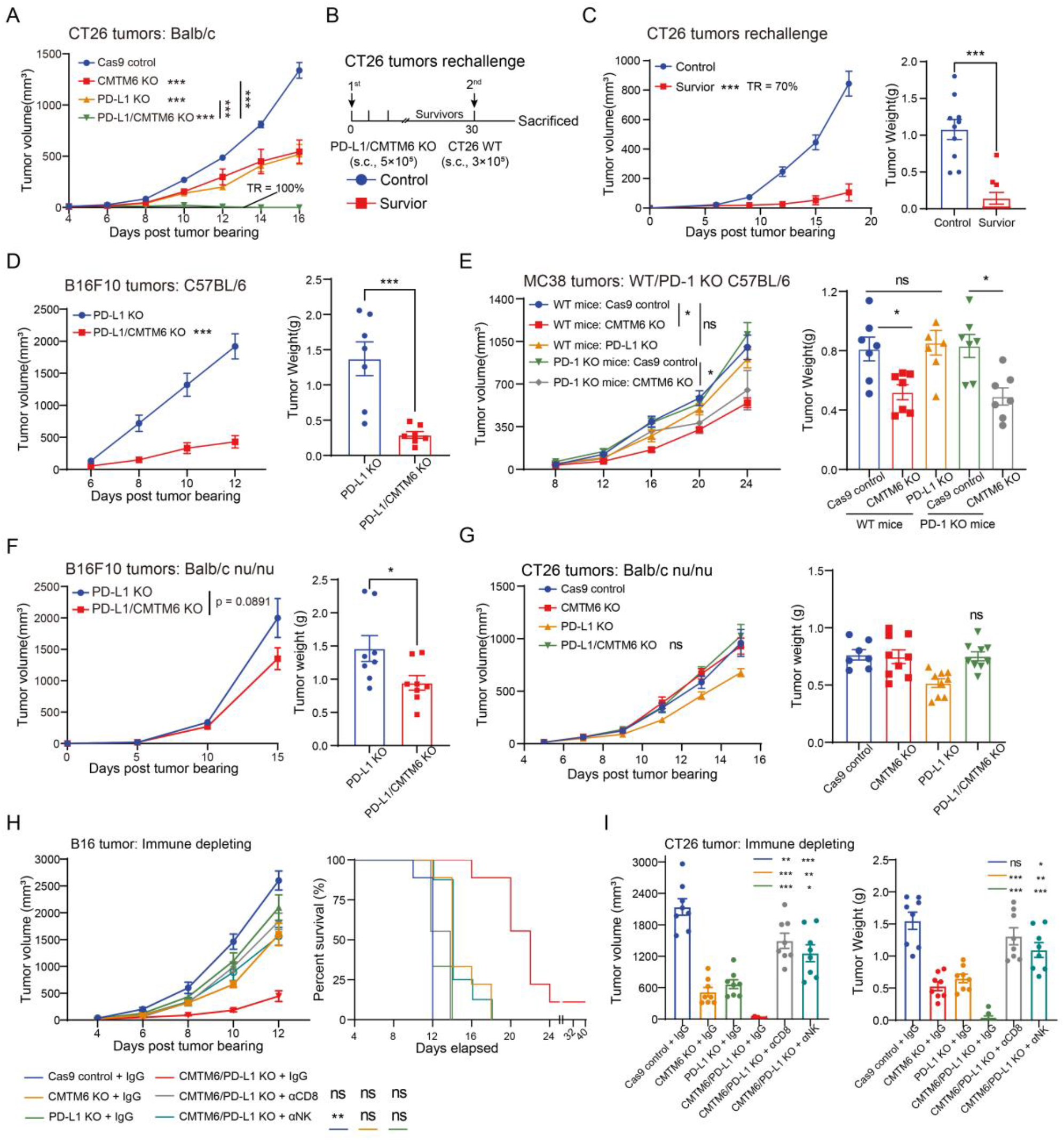
CMTM6 could participate in tumor immunity independently of the PD-1/PD-L1 axis. (**A**) Tumor growth kinetics of Cas9 control versus CMTM6 KO versus PD-L1 KO versus PD-L1/CMTM6 KO CT26 cells in Balb/c mice (n = 10). TR, tumor regression. (**B**) Schematic of the rechallenge schedules of mice with regressed tumors. (**C**) Tumor volume and weight are shown (n = 10) for rechallenge study. (**D**) Tumor growth kinetics of PD-L1 KO versus PD-L1/CMTM6 KO B16F10 cells in C57BL/6 mice (n = 7). And tumor weights of mice in two groups are shown. (**E**) Tumor growth kinetics of Cas9 control versus PD-L1 KO MC38 cells in C57BL/6 mice or PD-1^-/-^ mice (n = 7). And tumor weights of mice in different groups are shown. (**F**) Tumor growth kinetics of PD-L1 KO versus PD-L1/CMTM6 KO B16F10 cells in Balb/c nu/nu mice (n = 8). And tumor weights of mice in two groups are shown. (**G**) Tumor growth kinetics of Cas9 control versus CMTM6 KO versus PD-L1 KO versus PD-L1/CMTM6 KO CT26 cells in Balb/c nu/nu mice (n = 7). And tumor weights of mice in different groups are shown. (**H**) C57BL/6 mice bearing Cas9 control, CMTM6 KO, PD-L1 KO and PD-L1/CMTM6 KO B16F10 tumors dosed with immune cell deletion reagents. Tumor volume and survival curves are shown (n = 8). (**I**) Balb/c mice bearing Cas9 control, CMTM6 KO, PD-L1 KO and PD-L1/CMTM6 KO CT26 tumors dosed with immune cell deletion reagents. Tumor volume and weight are shown (n = 8). The data are presented as the mean ± SEM. * p < 0.05; ** p < 0.01; *** p < 0.001; ns not significant by unpaired t test or one-way ANOVA followed by Tukey’s multiple comparisons test.

Subsequently, in Balb/c nu/nu mice, PD-L1/CMTM6 KO was no longer identified as significantly inhibiting the growth of B16F10 and CT26 tumors compared to PD-L1 KO (Fig. 4, F and G). In addition, by deleting CD8^+^ T-cells and NK cells from mice, we found that in the case of PD-L1 deficiency, the tumor suppression effect of CMTM6 deletion remained mostly dependent on CD8^+^ T-cells and NK cells, especially CD8^+^ T-cells (Fig. 4, H and I).

The above results indicate that tumor CMTM6 can mediate tumorigenesis regulation independently of tumor PD-L1, and even the inhibitory effect of tumor CMTM6 ablation on tumor growth is largely unrelated to the PD-1/PD-L1 axis. The tumor immunomodulatory effects of CMTM6 separated from modulation of PD-L1 remain largely cytotoxic cell dependent.

### Host immune cell expression profile of CMTM6

Given that non-tumor cells may also express tumor-regulatory molecules, we further evaluated CMTM6 expression and function in host cells. Through searching the Human Protein Atlas database (*48*), we found that under normal physiological conditions, CMTM6 was widely expressed at the protein level in most human tissues (Fig. 5A), which was significantly different from the expression profile of PD-L1 that was only detected in few tissues such as the lung and colon (fig. S12A). At the single-cell level (*49, 50*), CMTM6 is preferentially expressed in granulocytes, monocytes, and macrophages both in immune organs and non-immune organs such as the liver and lung (fig. S11). And CMTM6 also exists in T-cells, B-cells, and NK cells (Fig 5B and fig. S11). In addition, CMTM6 is expressed abundantly in somatic cells, especially hepatocytes (fig. S11). Although PD-L1 is also highly expressed in granulocytes and monocytes/macrophages, its overall single-cell expression profile is significantly distinguished from that of CMTM6, and its expression level in the resting state is much lower than that of CMTM6 (fig. S12). A similar phenomenon was confirmed in single-cell sequencing data of mouse immune cells (Fig 5C). We then analyzed mouse splenocytes and human PBMCs by flow cytometry and found that CMTM6 was indeed broadly expressed in immune cell subsets at the protein level (Fig 5, D and E).

**Fig. 5.**
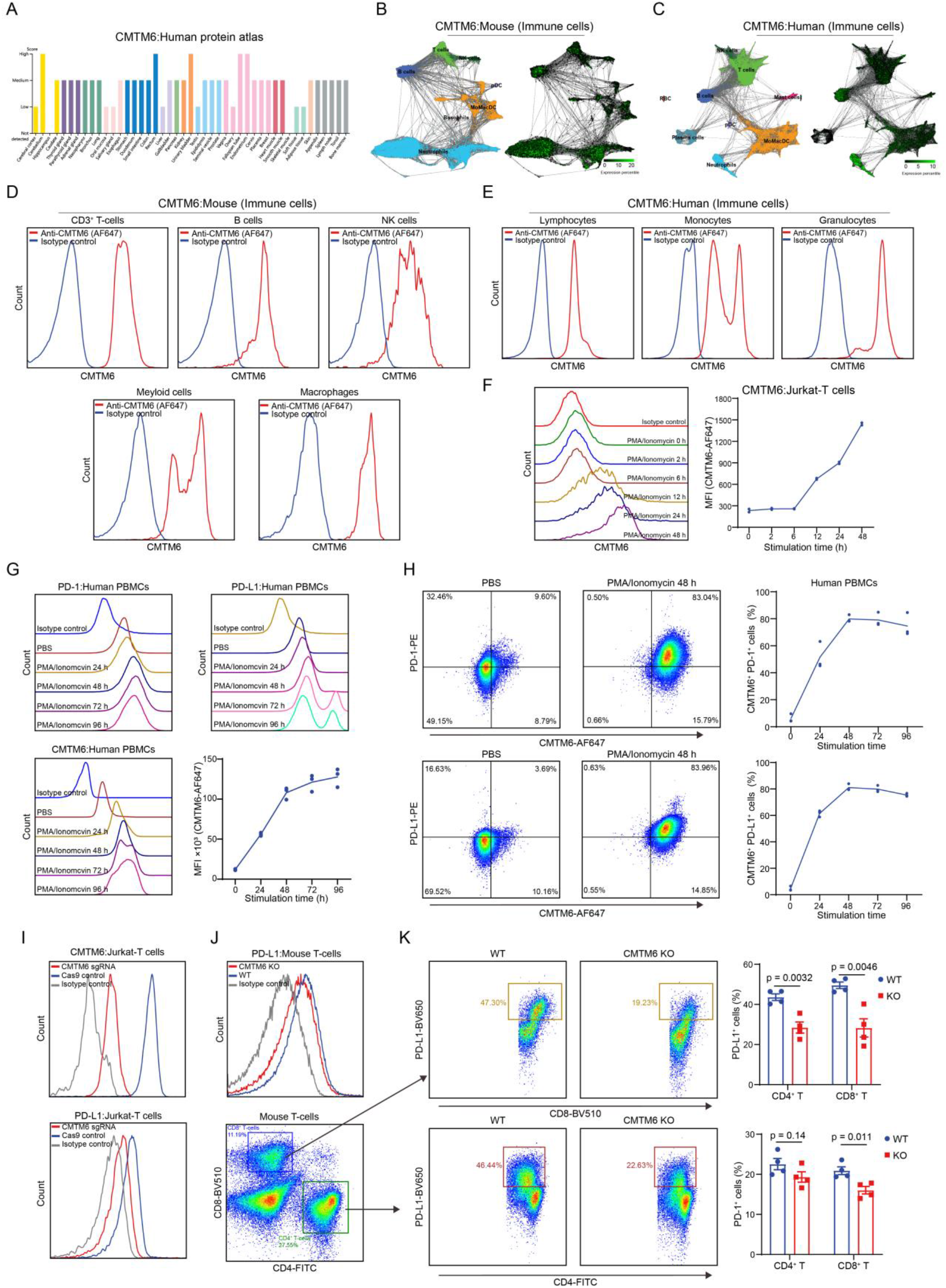
T-cells-intrinsic-CMTM6 regulated PD-L1 expression. (**A**) At the protein level, the expression of human CMTM6 in tissues of the whole body was analyzed by the Human Protein Atlas Project. (**B** and **C**) At the single-cell transcription level, the expression of human and murine CMTM6 in immune cells was analyzed. (**D**) Histograms show CMTM6 expression in different mouse spleen immune cells measured by flow cytometry (n = 3). (**E**) Histograms show CMTM6 expression in different human PBMC subsets by flow cytometry (n = 3). (**F**) Histograms and line plot show CMTM6 expression in Jurkat cells stimulated with PMA and ionomycin for different times measured by flow cytometry (n = 3). (**G**) Histograms and line plot show CMTM6/PD-1/PD-L1 expression in human PBMCs stimulated with PMA and ionomycin for different time measured by flow cytometry (n = 3). (**H**) Density plots and line plots show co-expression of CMTM6/PD-1 and CMTM6/PD-L1 in human PBMCs stimulated with PMA and ionomycin for different times measured by flow cytometry (n = 3). (**I**) Above shows the identification of CMTM6 knockout Jurkat cells. Below shows the difference in PD-L1 expression between Cas9 control and CMTM6 KO Jurkat cells stimulated by PMA and ionomycin. (**J**) The histogram shows the difference in PD-L1 expression between WT and CMTM6 KO murine CD3^+^ T-cells stimulated by PMA and ionomycin. (**K**) Density plots and scatter plots show the difference in PD-L1 and PD-1 expression between WT and CMTM6 KO murine CD4^+^ T-cells and CD8^+^ T-cells stimulated by PMA and ionomycin (n = 4). The data are presented as the mean ± SEM and analyzed by two-way ANOVA followed by Tukey’s multiple comparisons test.

### CMTM6 expression changed dynamically with T cell activation

Considering that the T-cell-intrinsic PD-L1 was recently found to play a non-negligible role in tumor progression, we selected the T-cell CMTM6 as the main research object to explore the association between CMTM6 and PD-L1 on host T-cells. First, in the PMA and ionomycin-activated Jurkat human T-cell model, CMTM6 expression rose progressively during the stimulation (Fig 5F). Following 48 hours of stimulation, Jurkat’s CMTM6 levels were approximately three times higher than in the resting state. Similarly, upon stimulation of human PBMCs with PMA and ionomycin, CMTM6 expression on CD3^+^ T-cells continued to increase, with a plateau obtained at around 96 hours (Fig 5G). PD-1 and PD-L1 expression on T-cells were also accompanied by increased activation time. Notably, co-expression of PD-1 and CMTM6 as well as PD-L1 and CMTM6 increased gradually with increasing activation time (Fig 5H). After 48 hours of stimulation, more than 80% of T-cells co-expressed CMTM6 and PD molecules.

### CMTM6 deficiency affected the membranal PD-L1 expression on T-cells

To access whether suppression of T-cell CMTM6 would also downregulate the membranal PD-L1 on T-cells, CMTM6-deficient Jurkat cells were constructed via the CRISPR-Cas9 system (Fig 5I). Membranal PD-L1 was significantly reduced in CMTM6 KO Jurkat cells compared to Cas9 control Jurkat cells 24 h after PMA and ionomycin stimulation (Fig 5I). Further, we constructed CMTM6^-/-^ mice and isolated CD3^+^ T-cells from their spleen and stimulated them with PMA and ionomycin for 24 h. Compared with splenic T-cells from WT mice, lower PD-L1 was found in CMTM6 KO splenic T-cells (Fig 5J). Specifically, CMTM6 deficiency reduced PD-L1 expression in activated mouse CD4^+^ T and CD8^+^ T-cells by 34.7% and 42.9%, respectively (Fig 5K). Moreover, CMTM6 deficiency did not affect PD-1 expression of activated CD4^+^ T and CD8^+^ T-cells. Hence, T-cell-intrinsic CMTM6 can also specifically affect the stability of PD-L1 on T-cell membranes.

### CMTM6 deficiency and T-cell development and activation

Besides, by analyzing the proportion of CD4^-^ CD8^-^ T-cells, CD4^-^ CD8^+^ T-cells, CD4^+^ CD8^+^ T-cells, and CD4^+^ CD8^-^ T-cells in the thymus of CMTM6-deficient mice and the co-expression of CD44 and CD25 in the double-negative (DN) T-cells, we preliminarily concluded that CMTM6 knockout had a slight impact on thymus T-cell development (fig. S13, A and B). Specifically, CMTM6 deficiency slightly raised the proportion of CD4^+^ CD8^-^ T-cells, slightly decreased the proportion of CD4^+^ CD8^+^ T-cells, and increased the proportion of DN4 cells. Furthermore, freshly isolated spleen T-cells from CMTM6-deficient mice also showed an increased preactivated phenotype indicated by CD69 and some other molecules (fig. S13C). However, the development and pre-activation phenotype of T-cells in the spleen of CMTM6-deficient mice did not differ from WT mice (fig. S13, D and E). Notably, CD4^+^ T-cells and CD8^+^ T-cells from the spleen of CMTM6-deficient mice were significantly higher in expression of CD44, CD69, IFN-γ and TNF-α than T-cells from WT mice after being isolated and stimulated with PMA and ionomycin for 24 hours, suggesting a higher activation state (fig. S14). These results suggest that CMTM6 has a potential regulatory effect on T-cell development and pre-activation in the thymus but does not affect T-cell development and pre-activation in the spleen. And CMTM6 may be associated with negative regulation of T-cell activation levels.

### The effect of host CMTM6 deficiency on tumor growth depended on CD8^+^ T-cells

To evaluate the effect of host CMTM6 deficiency on tumor development, we monitored the growth of B16F10 and MC38 tumors in CMTM6^-/-^, CMTM6^-/+^, and CMTM6^-/-^ mice (Fig 6A). The individual tumor growth kinetics and survival curves of B16F10 tumor-bearing mice showed that the higher the defective level of host CMTM6 the stronger inhibition for the tumor growth.

**Fig. 6.**
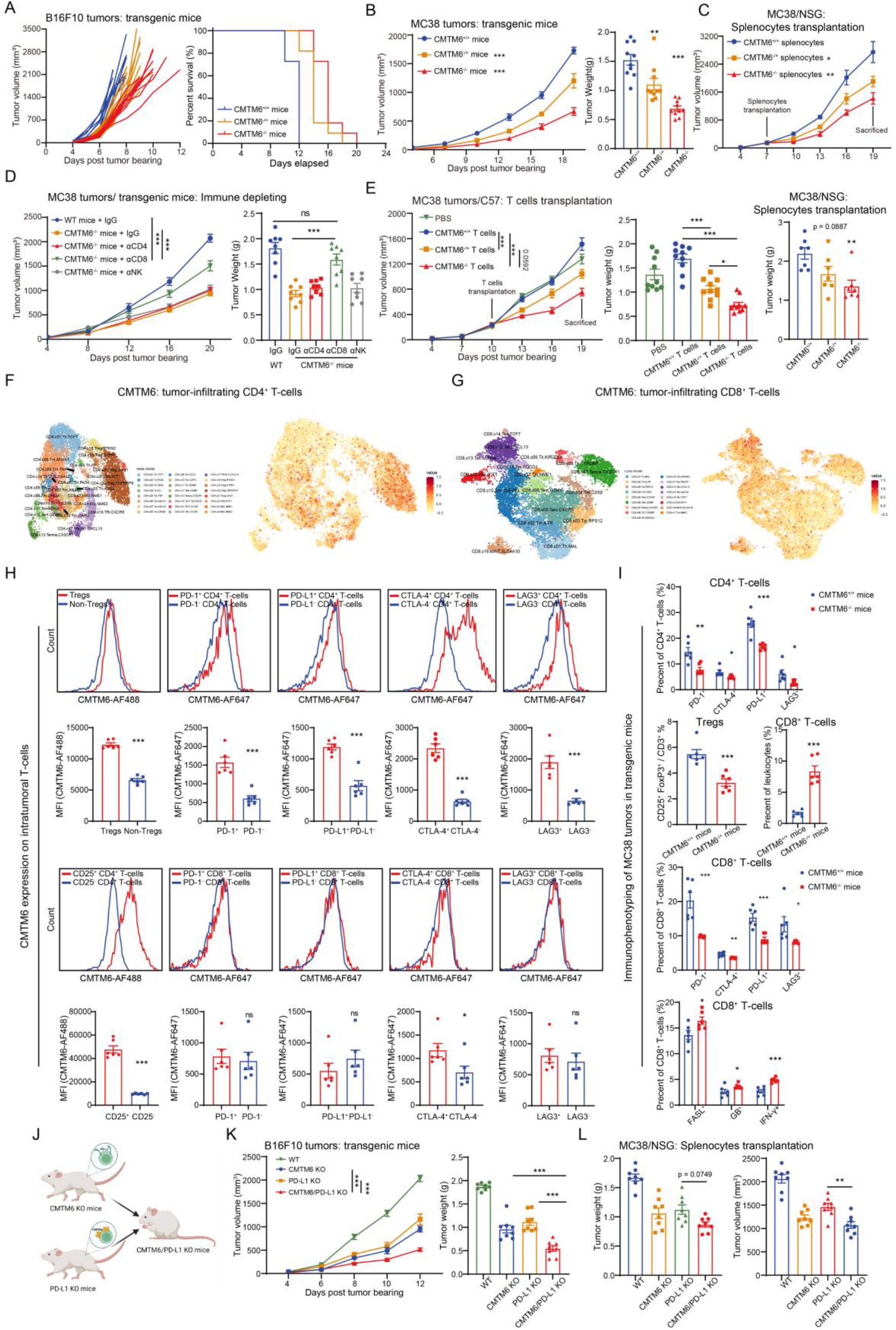
Host CMTM6 was associated with T-cell exhaustion and host CMTM6 knockout could enhance anti-tumor immunity independently of PD-L1. (**A**) Individual growth curves are shown for B16F10 tumors in CMTM6^+/+^, CMTM6^-/+^ and CMTM6^-/-^ mice (n = 8). Right is the Kaplan-Meier plot of survival. (**B**) Tumor growth kinetics of MC38 cells in CMTM6^+/+^, CMTM6^-/+^ and CMTM6^-/-^ mice (n = 10). And tumor weights of mice in three groups are shown. (**C**) Tumor growth kinetics of MC38 cells in NSG mice injected with CMTM6^+/+^, CMTM6^-/+^ and CMTM6^-/-^ splenocytes (n = 7). And tumor weights of mice in three groups are shown below. (**D**) WT and CMTM6^-/-^ mice bearing MC38 tumors dosed with immune cell deletion reagents. Tumor volume and weight are shown (n = 8). (**E**) Tumor growth kinetics of MC38 cells in C57BL/6 mice injected with CMTM6^+/+^, CMTM6^-/+^ and CMTM6^-/-^ T-cells (n = 10). And tumor weights of mice in four groups are shown. (**F** and **G**) Single cell RNA-seq data for CMTM6 expression in tumor-infiltrating CD4^+^ (F) and CD8^+^ (G) T-cells were analyzed. (**H**) Differences in CMTM6 expression in immune cells isolated from MC38 tumors in WT and CMTM6^-/-^ mice (n = 6). (**I**) At 19 days after tumor bearing, immune effector cells from MC38 tumors in WT and CMTM6^-/-^ mice were quantified by flow cytometry (n = 6). The percentages of specific CD4^+^ and CD8^+^ T-cells are shown based on their respective markers. (**J**) The schematic diagram shows the construction process of CMTM6/PD-L1 KO mice. (**K**) Tumor growth kinetics of B16F10 cells in WT, CMTM6 KO, PD-L1 KO and CMTM6/PD-L1 KO mice (n = 8). And tumor weights of mice in four groups are shown. (**L**) Tumor weight and volume of MC38 tumors in NSG mice injected with WT, CMTM6 KO, PD-L1 KO and CMTM6/PD-L1 KO splenocytes (n = 8) are shown. The data are presented as the mean ± SEM. * p < 0.05; ** p < 0.01; *** p < 0.001; ns not significant by unpaired t test or one-way/two-way ANOVA followed by Tukey’s multiple comparisons test.

Similarly, MC38 tumor development was significantly constrained in both CMTM6^-/+^ and CMTM6^-/-^ mice compared to MC38 tumors in WT mice, with an IR of 27.5% and 54.7%, respectively (Fig 6B).

To further investigate the effect of CMTM6 deficiency in host immune cells on their anti-tumor activity, we isolated splenocytes from CMTM6^-/-^, CMTM6^-/+^, and CMTM6^-/-^ mice and transfused them back into MC38 tumors of NSG mice (Fig 6C). The tumor growth curves showed that significant discrepancies in growth kinetics occurred between the three groups after transfusion. Splenocytes from CMTM6^-/-^ mice inhibited MC38 tumors most effectively, suggesting that CMTM6 deficiency enhanced the anti-tumor activity of the immune system.

Then, MC38 tumor-bearing CMTM6^-/-^ mice were given immune cell-deleting antibodies to remove intratumoral CD4^+^ T-cells, CD8^+^ T-cells and NK cells. The results showed that the intratumoral infiltration defect of CD8^+^ T-cells largely erased the tumor inhibitory effect in CMTM6-deficient mice (Fig 6D). Furthermore, infusion of splenic T-cells into MC38 tumors in C57BL/6 mice revealed that both CMTM6 partially and fully defective T-cells significantly limited tumor growth (Fig 6E).

The above results demonstrate that host CMTM6 also affects tumor growth, especially in immune cells represented by CD8^+^ T-cells. Suppression of CMTM6 can facilitate the anti-tumor activity of immune cells as the level of deficiency deepens.

### T-cell CMTM6 expression was associated with exhaustion phenotypes

Therefore, we speculated that CMTM6 was associated with tumor immunosuppression or exhaustion. By analyzing published pan-cancer tumor-infiltrating T-cell single cell sequencing data (*51*), we found a propensity for CMTM6 expression in Tregs as well as in exhausted CD8^+^ T (Tex) cells (Fig 6, F and G). Interestingly, we did not observe PD-L1 upregulation in CD8^+^ Tex, although we also noticed high expression of PD-L1 in Tregs (fig. S15). We then analyzed the CMTM6 expression profile of MC38 tumor-infiltrating WT T-cell lymphocytes by flow cytometry. We found higher levels of CMTM6 expression in Tregs, CD25^+^, PD-1^+^, PD-L1^+^, CTLA-4^+,^ and lymphocyte-activation gene 3 (LAG-3)^+^ T-cells (Fig 6H). And this co-expression with immunosuppressive molecules was more pronounced in CD4^+^ T-cells than in CD8^+^ T-cells. Further, we examined the proportion and function of immune cell subsets in MC38 tumors of CMTM6-deficient mice. The results showed a significant downregulation of CD4^+^ T and CD8^+^ T-cells expressing PD-1^+^, PD-L1^+^, CTLA-4^+^, LAG-3^+^, a decrease in Treg infiltration, an increase in CD8^+^ T-cell infiltration and a significant increase in FasL^+^, granzyme B^+^, IFN-γ^+^ CD8^+^ T-cells in MC38 tumors from CMTM6-deficient mice compared to that of WT mice (Fig 6I). These results suggest a link between CMTM6 and the exhaustion phenotype of tumor-infiltrating T-cells, and that defects in CMTM6 in host immune cells can lessen immunosuppression and augment cytotoxic effects.

### Host CMTM6 deficiency inhibited tumor growth in absence of the PD-1/PD-L1 axis

In PD-L1^-/-^ mice, we also aimed to investigate the biofunction of host CMTM6. We constructed CMTM6/PD-L1 double-knockout mice by breeding CMTM6 KO mice with PD-L1 KO mice (Fig 6J). Compared to PD-L1 KO mice and CMTM6 KO mice, B16F10 tumors in CMTM6/PD-L1 KO mice grew more slowly and had a lower tumor weight with an IR of 70.8% compared to WT mouse tumors (Fig 6K). Then we isolated splenocytes from WT, CMTM6 KO, PD-1 KO and CMTM6/PD-L1 KO mice and transfused them back into MC38 tumors of NSG mice. The CMTM6/PD-L1 KO splenocyte transfusion inhibited tumor growth more significantly than the PD-L1 KO splenocyte transfusion, suggesting that CMTM6 ablation from host immune cells could mediate anti-tumor immunity independent of PD-L1 (Fig 6L).

### scAAV-mediated CMTM6 suppression was effective in tumor immunotherapy

In accordance with our finding that ablation of tumor or host CMTM6 could promote anti-tumor responses, we have attempted to develop a self-complementary adeno-associated virus serotype 9 (scAAV9) encoding Cmtm6 mRNA targeting shRNA to mediate tumor gene therapy through CMTM6 suppression. Such a selection of serotypes and genomic structures can facilitate efficient infection and rapid expression in vivo. In vitro, the scAAV9-shCMTM6 effectively infected CT26 and B16F10 tumor cells and interfered with their CMTM6 expression (fig. S16).

In vivo, analysis of EGFP and CMTM6 expression within CT26 tumors three days after injection revealed that the scAAV9-shCMTM6 infected mainly tumor cells and only slightly infected immune cells (fig. S17).

CT26, B16F10 and Hepa1-6 tumor-bearing mice were administered with shNC scAAV9 and shCMTM6 scAAV9 on day 4, day 6 and day 8 (Fig 7A). In terms of growth kinetics and tumor weight, shCMTM6 scAAV9 successfully halted tumor progression with IRs of 48.9%, 58.8% and 50.6% against CT26, B16F10 and Hepa1-6 tumors, respectively (Fig 7, B to D). In addition, we generated a scAAV9 encoding both shCMTM6 and shPD-L1, and in vivo treatment experiments showed that shCMTM6&PD-L1 scAAV9 exerted superior antitumor efficacy over shCMTM6 scAAV9 and shPD-L1 scAAV9 with an IR of 76.4% (Fig 7, E and F). Further, immunophenotyping was performed to determine the proportion of immune cells in the tumor after scAAV9 treatment. The results showed that both shCMTM6 scAAV9 and shCMTM6&PD-L1 scAAV9 reduced PD-L1 levels in tumor cells and myeloid cells, decreased LAG-3, PD-1, PD-L1 expression in CD8^+^ T-cells, and promoted infiltrating CD8^+^ T-cells to express granzyme B, IFN-γ, perforin TNF-α (Fig 7G). Of course, shCMTM6 & PD-L1 scAAV9 had a more significant effect on TME.

**Fig. 7.**
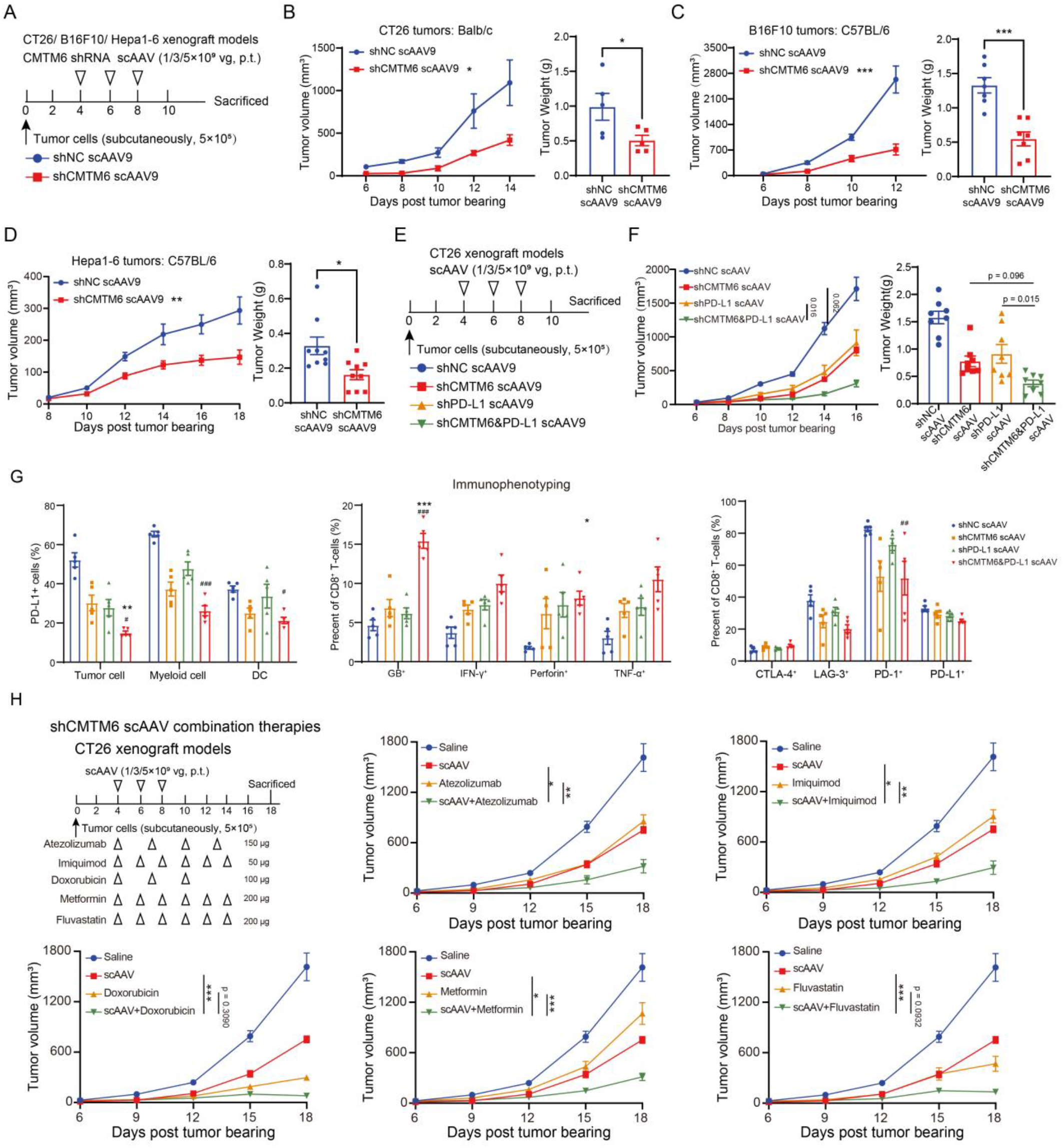
scAAV-mediated CMTM6 suppression was effective in tumor immunotherapy and combination therapy. (**A**) Schematic of treatment schedules and dosing of shNC and shCMTM6 scAAV9 against CT26, B16F10 and Hepa1-6 tumors. (**B** to **D**) Tumor growth kinetics of CT26, B16F10 and Hepa1-6 cells in mice treated with shNC or shCMTM6 scAAV9. And tumor weights of mice are shown. (**A**) Schematic of treatment schedules and dosing of shNC, shCMTM6, shPD-L1 and shCMTM6&PD-L1 scAAV9 against CT26 tumors. (**F**) Tumor growth kinetics of CT26 cells in mice treated with shNC, shCMTM6, shPD-L1 or shCMTM6&PD-L1 scAAV9 (n = 8). And tumor weights of mice are shown. (**G**) At 16 days after tumor bearing, immune effector cells from CT26 tumors in mice treated with shNC, shCMTM6, shPD-L1 or shCMTM6&PD-L1 scAAV9 were quantified by flow cytometry (n = 5). The percentages of specific CD4^+^ and CD8^+^ T-cells are shown based on their respective markers. (**H**) Schematic of combination treatment schedules against CT26 tumors is shown. Tumor growth kinetics of CT26 cells in mice treated with different combination strategies are shown (n = 8). The data are presented as the mean ± SEM. * p < 0.05; ** p < 0.01; *** p < 0.001; ns not significant by unpaired t test or one-way/two-way ANOVA followed by Tukey’s multiple comparisons test.

### scAAV-mediated CMTM6 suppression was effective in combination therapy

Then, we tried to combine shCMTM6 scAAV9 with anti-PD-L1 antibody Atezolizumab, TLR7 agonist imiquimod, chemotherapeutic drug Doxorubicin, antidiabetic drug Metformin and antilipemic drug Fluvastatin to further explore the value of shCMTM6 scAAV9 application (Fig 7H). These combination strategies represent the combination of immune checkpoint inhibitors, immune agonists, chemotherapeutic agents and metabolic modulating agents, respectively. In the tumor model of CT26, we found that all selected combinations achieved significantly better antitumor activity than single agents. Although none showed synergistic efficacy, the combinations did exert potent tumor suppressive effects, particularly shCMTM6 scAAV9 in combination with Doxorubicin and shCMTM6 in combination with Fluvastatin with IRs of 83.1% and 63.3%, respectively. The above results demonstrate that the shCMTM6 scAAV9 we developed can effectively mediate tumor immunotherapy and be applied in combination. CMTM6 is a novel tumor immunotherapy target with great clinical application potential.

## DISCUSSION

Based on studying of CMTM6 in regulating tumor development and immunity in this work, we developed AAV-mediated CMTM6-targeting gene therapy, confirming the potential of this previously unexploited target. As candidates, other gene therapies can also be wielded to target CMTM6, such as oncolytic viruses and siRNA-loaded-liposomes. In addition to gene therapy, classical antibodies and small-molecules are also expected to be screened for interfering with CMTM6 function. Since that CMTM6 is a four-transmembrane and its extracellular segment is predicted to be two little loops comprising only seven amino acids each, which makes anti-CMTM6 antibody development a spiny challenge. And distinguished from gene therapy, whether and how antibodies directly bound to CMTM6 can function still needs to be systematically evaluated. We postulated that affecting the CMTM6/PD-L1 interaction is one of the possible mechanisms for anti-CMTM6 antibodies. Of course, tailoring antibodies to link lysosomal-targeting peptides or ligands of cell surface lysosomal targeting receptor is promising direction to achieve direct degradation of CMTM6, which is consistent with the strategy of gene therapy. In addition, considering that the shCMTM6-scAAV9 we developed mainly infects tumor cells in vivo, modification of the AAV capsid with targeted peptides or antibodies (*52*) against T-cells and other immune cells is a potential way to broaden application and improve efficacy.

We found that the suppression of host/tumor CMTM6 contained tumor development incompletely dependent on PD-L1. In the MC38 tumor model, PD-L1 knockout did not affect tumor growth in vivo at all, whereas CMTM6 knockout could significantly repress tumors, which even suggests that CMTM6 inhibition can exert tumor-modulation completely independent of PD-L1. Thus, CMTM6 can function independently of PD-L1 as a novel tumor growth and immune regulating molecule. Of course, how CMTM6 functions independently of PD-L1 still needs to be explored. We hypothesized that CMTM6 might affect the stability of other immune-related membrane proteins. And as a MARVEL domain-containing membrane protein, CMTM6 may also be involved in regulating processes such as vesicle transport and tight junctions (11). In addition, some potential interaction molecules and functions of CMTM6 recently discovered (35-39) are also potential explanations. For example, CMTM6 may influence tumor immunogenicity by affecting tumor genome stability (37) and thus participate in tumor immunity. These findings warrant further investigation to ascertain the circumstances under which CMTM6 functions independently of PD-L1.

Considering that T-cells play a major antitumor role in CMTM6-deficient mice, we focused on the function of T-cell-intrinsic CMTM6. However, single-cell sequencing results showed that CMTM6 was widely expressed in other immune cells, and was in fact predominated in granulocytes and macrophages. For example, Markus Zeisbrich et al. found that CMTM6 in monocytes can affect the expression of membrane PD-L1 (*13*). Therefore, the function of CMTM6 on these immune cells is also worthy of investigation. And both the functions dependent of PD-L1 or not need to be systematically studied in other immune cells.

Of course, our present study still has some limitations, and we plan to address those issues in further research. First, since the in vivo experiments all used subcutaneous tumor models, and as in situ tumors can more realistically reflect the biofunctions, we plan to construct CMTM6-deficient in situ tumor models for further studies. Second, the evaluation of shCMTM6-scAAV9 is not yet complete. The efficacy needs to be evaluated in more tumor models and in situ tumor models, and the long-term efficacy and safety of this therapy needs to be paid attention to. Third, we plan to explore the molecules regulated by CMTM6 besides PD-L1, as well as the functions of CMTM6 on other immune cells to further substantiate the conclusions of this paper mechanistically.

In summary, based on the clinical relevance of CMTM6, this work analyzed and characterized CMTM6 expression profiles and the association between immune exhaustion phenotypes, and systematically investigated tumor/host CMTM6 in the regulation of tumor progression and tumor immunity in pan-cancer. Both host and tumor CMTM6 deficiency can reshape the anti-tumor immune microenvironment, which is mainly dependent on cytotoxic cells, in a PD-L1 non-dependent manner. And this study developed CMTM6-targeted gene therapy and demonstrated its potent pan-cancer efficacy for immunotherapy and manifold combinations. CMTM6 is a novel target with great potential for antitumor immunotherapy.

## MATERIALS AND METHODS

### Cells

Jurkat, RKO, CT26, B16F10, MC38, 4T1, LLC, Hepa1-6, and AAV-293 cell lines were purchased from Procell. B16-F10, LLC, Hepa1-6 and 293-AAV cells were cultured in Dulbecco’s Modified Eagle Medium (DMEM; Meilunbio) supplemented with 1% penicillin/streptomycin (P/S; Invitrogen) and 10% heat-inactivated fetal bovine serum (FBS; Life Technologies). Jurkat, CT26, MC38, and 4T1 were cultured in RPMI1640 medium (Meilunbio) containing 1% P/S and 10% FBS. RKO cells were cultured in Minimum Essential Medium (MEM; Meilunbio) supplemented with 1% P/S and 10% FBS. Other stable cell lines were constructed by the lentivirus system and cultured under the same conditions as the parental cells. Human PBMCs were purchased from Milestone. All cells were cultured at 37°C in a 5% CO _2_ humidified atmosphere.

### Plasmids and viruses

All the plasmids were purchased from Youbio. pVSV-G, pMDlg/pRRE, and pRSV-Rev were used to package lentiviruses. pLKO.1-TRC was used to express shRNA. LentiCRISPRv2 was used to express sgRNA. pscAAV-ZsGreen1-shRNA, pHelper, and pAAV-RC were used to package scAAV. shRNA sequences: mouse CMTM6, 5′-TGCCTAACAGAAAGCGTGT-3′; mouse PD-L1, 5′-CCGAAATGATACACAATTCGA-3′; non-targeting, 5′-GGGTA TCGACGATTACAAA-3′. sgRNA sequences: mouse CMTM6, 5′-CCTGGCCGCCTACTTCGTCC-3′; mouse PD-L1, 5′-GTATGGCAGCAACGTCACGA-3′; human CMTM6, 5′-CCGGGTCCTCCTCCGTAGTG-3′.

### Mice

Female (Six-to eight-week-old) BALB/c and C57BL/6 mice were purchased from the Shanghai Slack. CMTM6^-/-^, PD-L1^-/-^ and PD-1^-/-^ mice were purchased from GemPharmatech. CMTM6 and PD-L1 double knockout mice were obtained by CMTM6^-/-^ and PD-L1^-/-^ mice hybridization. All mice were maintained under specific pathogen-free (SPF) conditions in the animal facility of the Shanghai Institute of Materia Medica, Chinese Academy of Sciences (SIMM). Animal care and experiments were performed in accordance with the protocols approved by the Institutional Laboratory Animal Care and Use Committee (IACUC).

### TCGA cohorts analysis

The clinical pan-cancer CMTM6 mRNA expression data were obtained from the TCGA datasets (https://portal.gdc.cancer.gov/). Expression differences, prognostic correlation, and immunological correlation were analyzed by GEPIA software and R software v4.0.3.

### Tumor models of transgenic tumor cells or transgenic mice

For the subcutaneous model, the mice were randomly grouped and injected subcutaneously with 5×10 ^5^ of tumor cells in the right forelimb. Tumor volume was measured and recorded every two or three days. Tumor volume was calculated as tumor volume (mm^3^) = tumor length×width×width/2, and tumor growth curves were plotted. At least five time points after collection, the mice were sacrificed and the tumors harvested for weighing, photographing, or other purposes. For survival studies, euthanasia endpoints were set as tumors larger than 2000 mm^3^ or larger than 20 mm in length or mice with significant weight loss. For the lung metastasis model, 1×10 ^6^ of B16F10 cells were injected into C57BL/6 mice or Balb/c nu/nu mice through the tail vein. After 20 days, the mice were sacrificed and the lungs were harvested for weighing and photographing.

### In vivo treatment studies

Studies of CTLA-4 antibody and PD-L1 antibody therapy in 4T1 and LLC tumors, as well as studies of scAAV monotherapy and combination treatment, are consistent with the descriptions above. The administration methods of each drug are shown in the flow chart in Fig 2 and Fig 7.

### Immune cell depletion study

Mice were randomly divided and injected with tumor cells subcutaneously, which was defined as day 0. CD4^+^ T-cells were deleted by intraperitoneal injection of anti-CD8 (BioXcell) antibodies on day -2 (150 μg/each), day 0 (150 μg/each), day 4 (150 μg/each) and day 8 (150 μg/each).

CD8^+^ T-cells were deleted by intraperitoneal injection of anti-CD8β (BioXcell) antibodies on day -2 (100 μg/each), day 0 (100 μg/each), day 4 (100 μg/each) and day 8 (100 μg/each). NK cells of C57BL/6 mice were deleted by intraperitoneal injection of anti-NK1.1 (BioXcell) antibodies on day -2 (200 μg/each), day 0 (200 μg/each), day 4 (100 μg/each) and day 8 (100 μg/each). NK cells of Balb/c mice were deleted by intraperitoneal injection of anti-Asialo-GM1 (Biolegend) antibodies on day -2 (35 μL/each), day 0 (35 μL/each), day 4 (35 μL/each) and day 8 (35 μL/each). Macrophages were deleted by chlorophosphate-liposomes (200 μL/each/ intraperitoneally, Yeasen) on day -2, 4 day 8. CD3^+^ T-cells were deleted by FTY720 (Meilunbio) on day -1 (30 μg/each), day 0 (30 μg/each) and day 1 to 12 (6 μg/each). The deletion effect was identified by flow cytometry.

### Immunophenotype analysis

For immunotyping of intratumoral cells, tumor tissues were digested by collagenase IV (Yeasen) and hyaluronidase (Yeasen) into single cell suspensions and then were subjected to cell extraction by lymphocyte separation medium (Dakewei) for lymphocyte sorting or were subjected to erythrocyte removal by red blood cell lysis buffer (Yeasen) for non-lymphocyte staining. Then the cells were blocked with 4% FBS and anti-CD16/CD32 (BD Biosciences), incubated with surface marker antibodies for 20 minutes at 4° C and then permeabilized with BD Cytofix/Cytoperm buffer before intracellular labeling antibodies were added for 30 minutes at 4° C. Then transcription factors were labeled for 45 minutes at 4° C by antibodies after permeabilized with BD TF Fix/Perm buffer. Flow cytometry analysis was performed using ACEA NovoCyte and data processing was done through NovoExpress software. Antibody staining was performed following the manufacturer’s recommendations. For immunotyping of splenocytes and thymocytes, single-cell suspensions were obtained by grinding tissues and were subject to erythrocyte removal. The other steps are the same as above.

### In vitro cell assays

For analysis of dynamic changes in CMTM6 expression during activation of Jurkat cells, Jurkat cells were seeded in 24-well plates and stimulated with 50 ng/mL of PMA (Meilunbio) and 1 μg/mL of ionomycin (Meilunbio) for 0 h, 2 h, 6 h, 12 h, 24 h, and 48 h, respectively. After fixation and permeabilization, the cells were stained with anti-CMTM6 antibody (Abcam) and goat anti-rabbit IgG H&L (Alexa Fluor 647) antibody (Abcam) and detected by flow cytometry.

For the analysis of CMTM6 affecting Jurkat PD-L1, CMTM6-deficient Jurkat cells were constructed through the CRISPR/Cas9 system. PD-L1 levels were detected by flow cytometry 24 h after Jurkat cells were stimulated with 50 ng/mL of PMA and 1 μg/mL of ionomycin.

For the cell proliferation assay, 1000 tumor cells were seeded in 96-well plates. CCK8 (Yeasen) was added at 0 h, 24 h, 48 h, 72 h, respectively, and the signal was read at OD 450 nm.

### Ex vivo cell assays

For the TILs-CT26 co-culture study, 3×10 ^5^ of CT26 cells were seeded in 24-well plates 12 h in advance. TILs were added to the wells at a ratio of 5:1 and 10:1, respectively. After 24 hours, TILs were removed and the supernatant was used for cytokine detection by ELISAs. The survival of CT26 cells was analyzed by CCK8 and annexin V-FITC/PI apoptosis detection kit (Yeasen).

The C57BL/6 mice were sacrificed, and the spleens and thymuses were collected for single-cell suspensions. Without stimulation, splenocytes and thymocytes were detected for pre-activated phenotypes and cell development by flow cytometry using specific antibody staining. 2×10 ^6^ of splenocytes were added to 12-well plates and were stimulated with 50 ng/mL of PMA and 1 μg/mL of ionomycin for 24 h for activation phenotype analysis or PD-L1 expression analysis, or for continuous timepoints for dynamic change of CMTM6 expression analysis.

Freshly recovered human PBMCs (Milestone) were seeded at 5×10 ^5^ of per well in 24-well plates. Without stimulation, CMTM6 expression of immune cell subsets in PBMCs was detected by flow cytometry. PBMCs were stimulated with 50 ng/mL of PMA and 1 μg/mL of ionomycin for 0 h, 24 h, 48 h, 72 h and 96 h. Subsequently, cell surface expression of PD-1, PD-L1 and CMTM6 was measured by flow cytometry.

### RKO/PBMCs humanized tumor model

Female NSG mice were randomly divided into groups of 8. PBMC cells (Milestone) derived from the same individual were revived and used immediately. Mice were inoculated subcutaneously with RKO cells (5×10 ^6^) mixed with human PBMCs (1×10 ^6^) or RKO cells only on the right flank. Tumor volumes were monitored every five days for five time points.

### Adoptive transfusion of immune cells

For adoptive transfusion of TILs, the donor mice were tumor-bearing six days before the recipient mice. On day 6 of tumor growth in recipient mice, the donor mice were sacrificed and tumor tissue was obtained. Under aseptic conditions, tumor tissues were digested by collagenase IV (Yeasen) and hyaluronidase (Yeasen) into single cell suspensions and then were subjected to cell extraction by lymphocyte separation medium (Dakewei) for lymphocyte sorting. TILs (5×10 ^6^) were peritumoral injected into recipient mice. For splenocyte and T-cell transfusion, under aseptic conditions, the spleen was processed as described above and the T-cells were obtained by magnetic bead sorting (Pan T Cell Isolation Kit II; Miltenyi). Splenocytes or T-cells (5×10 ^6^) peritumorally injected into recipient mice with tumor growth for 7 or 10 days.

### Analysis of scAAV infection

In vitro, 1×10 ^5^ of CT26 and B16F10 cells were inoculated in 24-well plates, respectively. After eight hours, scAAV was added to the well plates at a MOI of 10,000. After 72 hours of infection, the green fluorescent signal was detected by flow cytometry and fluorescence microscopy. In vivo, 5×10 ^9^ of scAAV were peritumorally injected at day 6 of tumor growth. After 3 days, the tumor tissues were collected and processed into single cell suspensions, and the green fluorescence signal in each cell subpopulation was analyzed by immunotyping.

### Single-cell sequencing data analysis

Single cell RNA-seq data used in this study were all from publicly available data. Data for CMTM6 expression in immune cells were obtained from the GEO database (GSE127465) and the Human Protein Atlas Project (https://www.proteinatlas.org/). The SpringViewer interactive tool was used to analyze the data. Data for CMTM6 expression in tumor-infiltrating T-cells were obtained from the GEO database (GSE156728) and analyzed by the scDVA interactive tool (http://cancer-pku.cn:3838/PanC_T/).

### RNA-Seq

RNA was isolated from fresh tumor tissues. Transcriptome libraries were constructed and sequenced by BGI. Differential expression was evaluated with DESeq. A fold-change of 2:1 or greater and a false discovery rate (FDR)-corrected p-value of 0.05 or less were set as the threshold for differential genes. Immune signature scores are defined as the mean log_2_(fold-change) among all genes in each gene signature. Cell infiltration within tumor tissues was estimated by xCell.

### Statistical analysis

The in vivo experiments were randomized but the researchers were not blinded to allocation during experiments and outcome analysis. Statistical analysis was performed using GraphPad Prism 8 Software. A Student’s t test was used for comparison between the two groups. Multiple comparisons were performed using one-way ANOVA followed by Tukey’s multiple comparisons test or two-way ANOVA followed by Tukey’s multiple comparisons test. Survival analysis was performed by the Kaplan-Meier method. Detailed statistical methods and sample sizes in the experiments are described in each figure legend. No statistical methods were used to predetermine the sample size. All statistical tests were two-sided and P-values < 0.05 were considered to be significant. ns not significant; *p < 0.05; **p < 0.01; ***p < 0.001.

## Supplementary Materials

Supplementary materials and methods

figs. S1 to S17

## Acknowledgments

We thank Professor Y.Geng for scientific discussions. All the figures were created with BioRender.com.

## Author contributions

L.G. and Y.L. designed the experiments and analyzed the data. Y.L. performed the experiments and prepared the manuscript. R.C. assisted in performing the experiments and preparing the manuscript. Y.T., X.P., F.L. and C.H. assisted in performing the animal experiments. X.Y. and J.S. assisted in data interpretation.

## Data and materials availability

All data are available in the main text or the supplementary materials. Correspondence and requests for materials should be addressed to L.G.

