## Supplementary Materials for "Suppression of tumor/host intrinsic CMTM6 drives anti-tumor cytotoxicity in a PD-L1 independent manner"

**This file contains:**

Supplementary materials and methods

Fig. S1 The association of CMTM6 expression with the prognosis in TCGA cohorts.

Fig. S2 Knockout of CMTM6 and its effect on tumor-cell growth in vitro.

Fig. S3 Regulation of PD-L1 via CMTM6 depletion in different mouse tumors.

Fig. S4 The association of CMTM6 expression with clinical tumor progression in TCGA cohorts.

Fig. S5 CMTM6-deficient RKO cells and the association of CMTM6 expression with immune infiltration and checkpoints expression in TCGA cohorts.

Fig. S6 Depletion of CMTM6 enhanced specific antitumor cytotoxicity.

Fig. S7 Flow-cytometry gating strategy for quantifying lymphocyte subsets in murine tumors.

Fig. S8 Flow-cytometry gating strategy for quantifying myeloid cells in murine tumors.

Fig. S9 Tumor CMTM6 knockout affected intratumoral adaptive immune responses, antigen processing, interferon process and NK cell functions.

Fig. S10 Knockout of CMTM6 and PD-L1 and its effect on tumor-cell growth in vitro.

Fig. S11 Expression profile of CMTM6 at the single cell level.

Fig. S12 Expression profile of PD-L1 at the single cell level.

Fig. S13 CMTM6 affected T cell development and pre-activation.

Fig. S14 CMTM6 affected T cell activation.

Fig. S15 Pan-cancer T-cell expression profile of PD-L1.

Fig. S16 scAAV effectively infected mouse tumor cells in vitro.

Fig. S17 scAAV effectively infected mouse tumor cells in vivo.

### Supplementary materials and methods

### Key reagents

| REAGENT or RESOURCE | SOURCE | IDENTIFIER |
|---|---|---|
| **Antibodies** | | |
| FITC Mouse Anti-Human CD3 | BD Bioscience | Cat# 561806 |
| Hu CD279 (PD-1) PE EH12.1 | BD Bioscience | Cat# 560795 |
| FITC Hamster Anti-Mouse CD3e | BD Bioscience | Cat# 553061 |
| APC Rat Anti-Mouse CD8a | BD Bioscience | Cat# 553035 |
| Ms CD16/CD32 Pure 2.4G2 | BD Bioscience | Cat# 553141 |
| Hu CD274 PE MIH1 | BD Bioscience | Cat# 557924 |
| BV786 Rat Anti-Mouse IFN-γXMG1.2 | BD Bioscience | Cat# 563773 |
| BV786 Rat Anti-Mouse CD274 MIH5 | BD Bioscience | Cat# 741014 |
| PE Mouse Anti-Human LAG-3 T47-530 | BD Bioscience | Cat# 565617 |
| Ms CD4 BV421 GK1.5 | BD Bioscience | Cat# 562891 |
| Ms CD4 FITC RM4-5 | BD Bioscience | Cat# 53046 |
| Ms CD274 BV650 MIH5 | BD Bioscience | Cat# 740614 |
| Ms LY-6G BV605 1A8 | BD Bioscience | Cat# 563005 |
| Ms I-A/I-E BV510 M5/114.15.2 | BD Bioscience | Cat# 742893 |
| Ms CD86 APC GL1 | BD Bioscience | Cat# 558703 |
| Ms CD3e APC-Cy7 145-2C11 | BD Bioscience | Cat# 557596 |
| Ms CD8a BV510 53-6.7 | BD Bioscience | Cat# 563068 |
| Ms CD11c BV786 HL3 | BD Bioscience | Cat# 563735 |
| Ms CD45.2 APC-Cy7 104 | BD Bioscience | Cat# 560694 |
| Ms CD45R/B220 PerCP-Cy5.5 RA3-6B2 | BD Bioscience | Cat# 552771 |
| Ms CD80 PE 16-10A1 | BD Bioscience | Cat# 553769 |

| | | |
|---|---|---|
| Ms TNF BV650 MP6-XT22 | BD Bioscience | Cat# 563943 |
| Ms CD279 BV786 J43 | BD Bioscience | Cat# 744548 |
| Ms CD178 BV421 MFL3 | BD Bioscience | Cat# 740054 |
| Ms CD49b/Pan-NK Cells FITC DX5 | BD Bioscience | Cat# 553857 |
| Ms CD62L BV650 MEL-1 | BD Bioscience | Cat# 564108 |
| Ms IL-17A BV605 TC11-18H10 | BD Bioscience | Cat# 564169 |
| Ms CD8a BV510 53-6.7 | BD Bioscience | Cat# 563068 |
| Ms CD44 BV421 IM7 | BD Bioscience | Cat# 563970 |
| Ms IL-4 PE-Cy7 11B11 | BD Bioscience | Cat# 560699 |
| Ms CD4 PerCP-Cy5.5 RM4-5 | BD Bioscience | Cat# 550954 |
| Ms CD25 PE 3C7 | BD Bioscience | Cat# 553075 |
| Ms I-A/I-E FITC 2G9 | BD Bioscience | Cat# 553623 |
| Brilliant Violet 421™ anti-mouse/human CD11b | BioLegend | Cat# 101235 |
| PE anti-mouse LAP (TGF-β1) | BioLegend | Cat# 106323 |
| Brilliant Violet 605™ anti-mouse CD152 | BioLegend | Cat# 106323 |
| Ultra-LEAF™ Purified anti-Asialo-GM1 Antibody | BioLegend | Cat# 146002 |
| PE-Cyanine7 Rat Anti-Mouse F4/80 | eBioscience | Cat# 25-4801-82 |
| ANTI-MO CD206 MR6F3 AF488 | eBioscience | Cat# 53-2061-82 |
| ANTI-MOUSE/RAT FOXP3 (FJK-16S) APC | eBioscience | Cat# 17-5773-82 |
| ANTI-MOUSE GRANZYME B (NGZB) PE-CYANINE | eBioscience | Cat# 25-8898-82 |
| ANTI-MO PERFORIN (EBIOOMAK-D) APC | eBioscience | Cat# 17-9392-80 |
| Anti-CMTM6 antibody [EPR23015-45] | Abcam | Cat# ab264067 |

| | | |
|---|---|---|
| Anti-beta Tubulin antibody [1E1-E8-H4] | Abcam | Cat# ab131205 |
| Goat Anti-Rabbit IgG H&L (Alexa Fluor® 488) | Abcam | Cat# ab150077 |
| Goat Anti-Rabbit IgG H&L (Alexa Fluor® 647) | Abcam | Cat# ab150079 |
| Goat Anti-Rabbit IgG H&L (HRP) | Abcam | Cat# Ab6721 |
| InVivoMab anti-mouse CTLA-4 (CD152) | BioXcell | Cat# BE0032 |
| InVivoMab anti-mouse CD4 | BioXcell | Cat# BE0119 |
| InVivoMab anti-mouse NK1.1 | BioXcell | Cat# BE0036 |
| InVivoMab anti-mouse CD8β (Lyt 3.2) | BioXcell | Cat# BE0223 |
| Human IgG1 | This lab | N/A |
| Atezolizumab | This lab | N/A |
| **Chemicals and recombinant proteins** | | |
| Native Human IgG1 protein | Abcam | Cat# ab90283 |
| Fingolimod HCL/FTY720 | Meilunbio | Cat# MB1552 |
| Doxorubicin HCl | Meilunbio | Cat# MB1087 |
| Fluvastatin sodium salt | Meilunbio | Cat# MB1420 |
| Phorbol 12-myristate 13-acetate (PMA) | Meilunbio | Cat# MB5349 |
| Ionomycin | Meilunbio | Cat# MB7511 |
| Mouse IFN-γ | Genscript | Cat# Z02916 |
| Human IFN-γ | Genscript | Cat# Z02986 |
| Brefeldin A (BFA) Solution | BD Pharmingen | Cat# 347688 |
| Atezolizumab | This lab | N/A |
| Imiquimod | MCE | Cat# HY-B0180 |
| Metformin | MCE | Cat# HY-B0627 |

| | | |
|---|---|---|
| Clodronate Liposomes | Yeasen | Cat# 40337ES08 |
| **Critical commercial kits** | | |
| Annexin V-FITC/PI Apoptosis Detection Kit | Yeasen | Cat# 40302ES20 |
| Cell Counting Kit（CCK-8） | Yeasen | Cat# 40203ES60 |
| Cytofix/Cytoperm Soln Kit | BD Pharmingen | Cat# 554714 |
| Transcription Factor Buffer Set | BD Pharmingen | Cat# 562574 |
| MOUSE IFN-γ ELISA KIT | ExCell | Cat# EM007-96 |
| MOUSE TNF-α ELISA KIT | ExCell | Cat# EM008-96 |
| Pan T Cell Isolation Kit II, mouse | Miltenyi | Cat# 130-095-130 |

### Supplementary figures

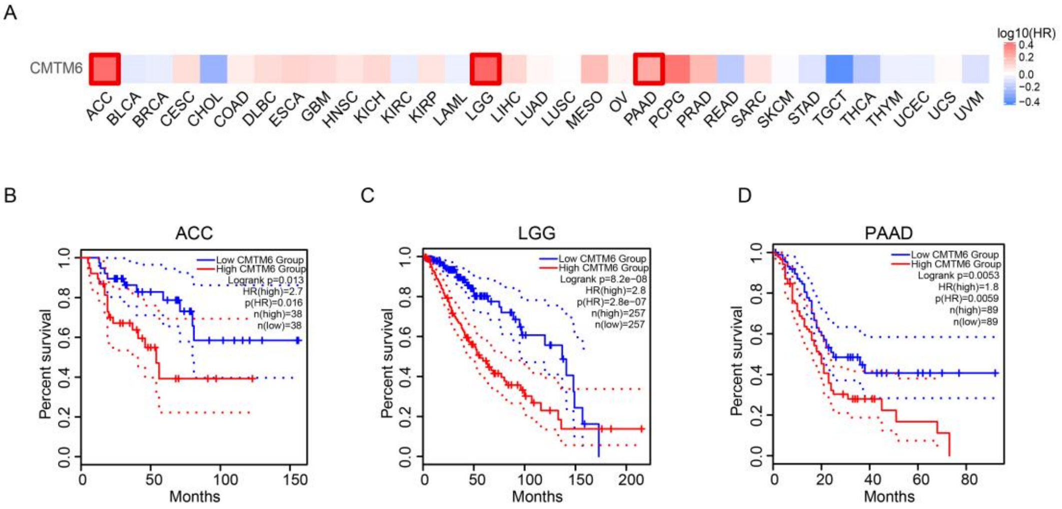


**Fig. S1. The association of CMTM6 expression with the prognosis in TCGA cohorts.** (**A**) The heat map shows the correlation between CMTM6 expression and patient survival with different tumor types as indicated in the TCGA database and determined by GEPIA. (**B** to **D**) The Kaplan-Meier curves show the overall survival in patients with adrenocortical carcinoma (ACC), low-grade glioma (LGG) or pancreatic adenocarcinoma (PAAD) from the TCGA database and determined by GEPIA.

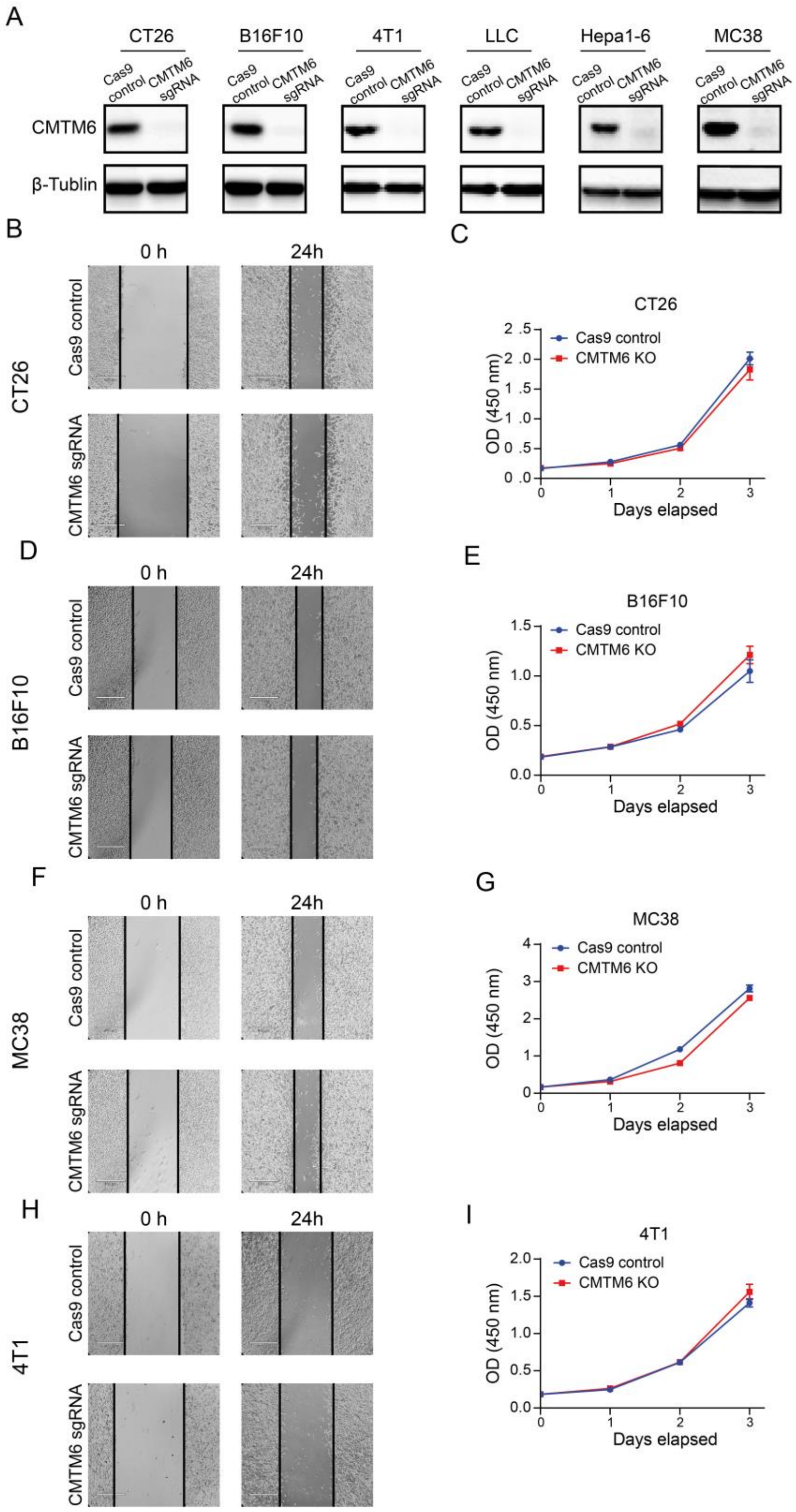


**Fig. S2. Knockout of CMTM6 and its effect on tumor-cell growth in vitro.** (**A**) Western blot analysis of CMTM6 expression in murine tumor lines with CMTM6 knockout. β-Tublin was used as the protein loading control. (**B** to **I**) The wound-healing assays (n = 3) and cell proliferation assays (n = 6) were performed on murine tumor lines with CMTM6 knockout. The data are presented as the mean ± SEM.

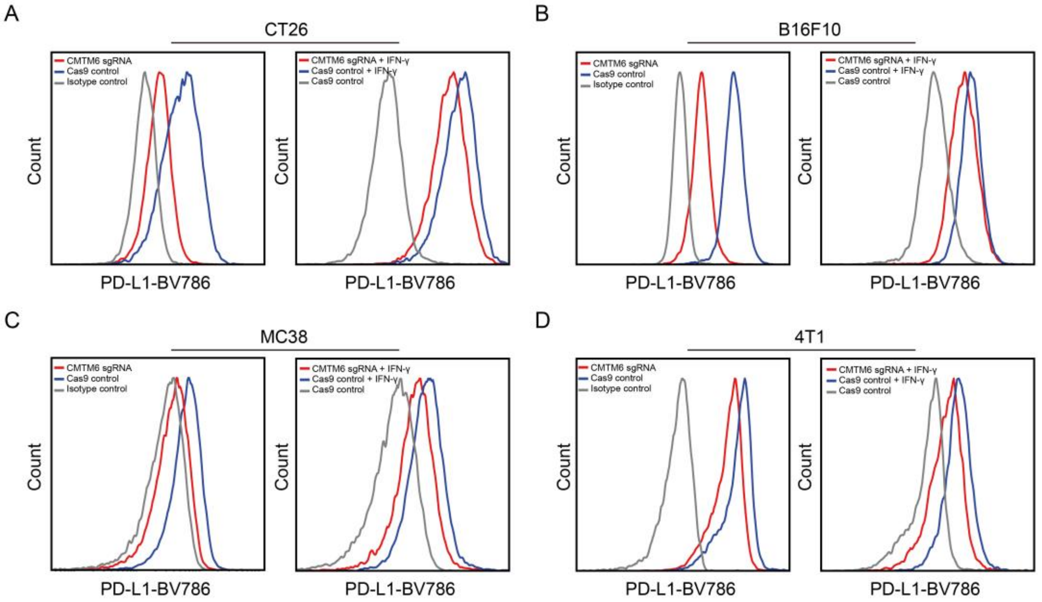


**Fig. S3. Regulation of PD-L1 via CMTM6 depletion in different mouse tumors.** (**A** to **D**) The histograms show the effect of CMTM6 deficiency on PD-L1 expression in CT26, B16F10, MC38 and 4T1 tumor cells with and without IFN-γ induction. The level of membranal PD-L1 was detected by flow cytometry.

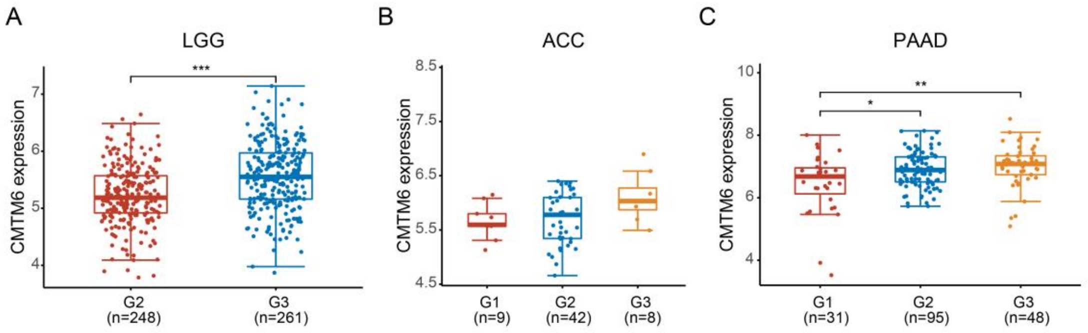


**Fig. S4. The association of CMTM6 expression with clinical tumor progression in TCGA cohorts.** (**A** to **C**) Among the three types of tumors (LGG, ACC, and PAAD) in which high CMTM6 indicated poor prognosis, CMTM6 mRNA expression was compared between different tumor grades provided by available TCGA data (missing G1 data in LGG). The data are presented as the mean ± SEM. * $p < 0.05$; ** $p < 0.01$; *** $p < 0.001$; ns not significant by unpaired t test or one-way ANOVA followed by Tukey’s multiple comparisons test.

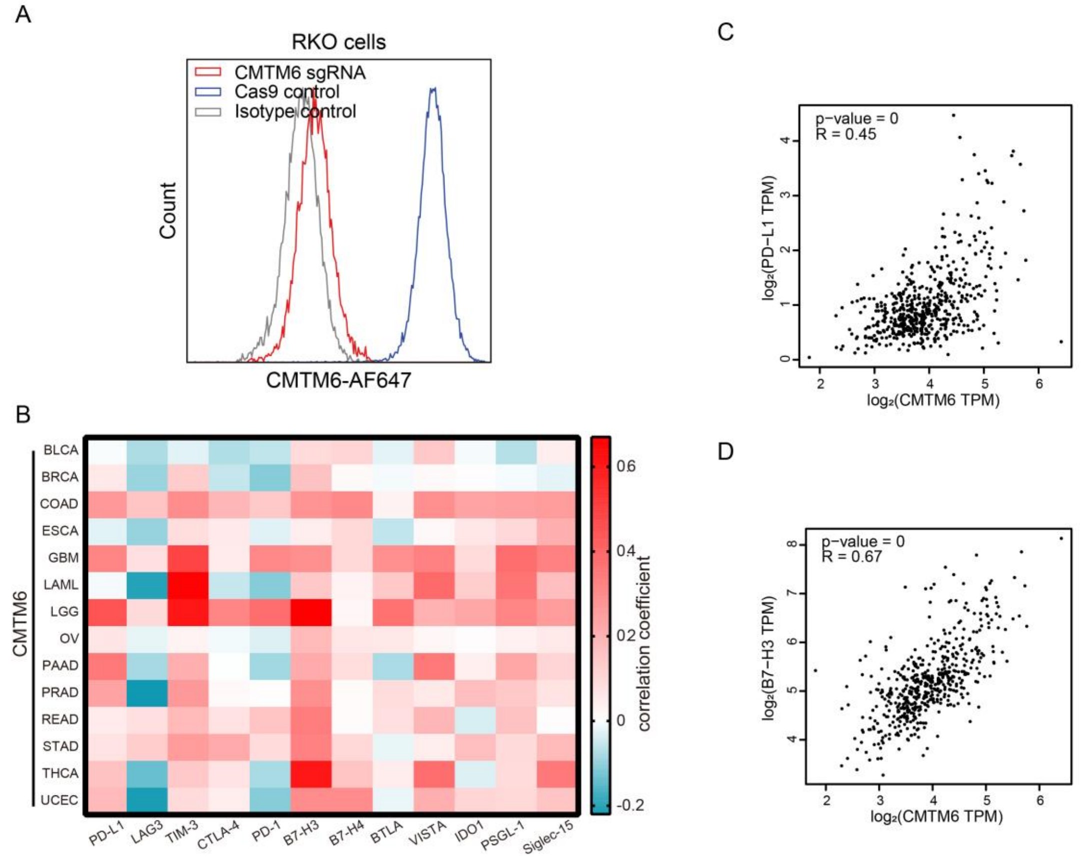


**Fig. S5. CMTM6-deficient RKO cells and the association of CMTM6 expression with immune infiltration and checkpoints expression in TCGA cohorts.** (**A**) Flow cytometry analysis of the expression of CMTM6 in RKO cells with CMTM6 knockout. (**B** to **D**) Correlation analysis between CMTM6 and several immune checkpoints in patients with different tumor types as indicated in the TCGA database and determined by GEPIA. The heat map shows the correlation coefficient in pan-cancer. Scatter plots are representative of the results of correlation analysis.

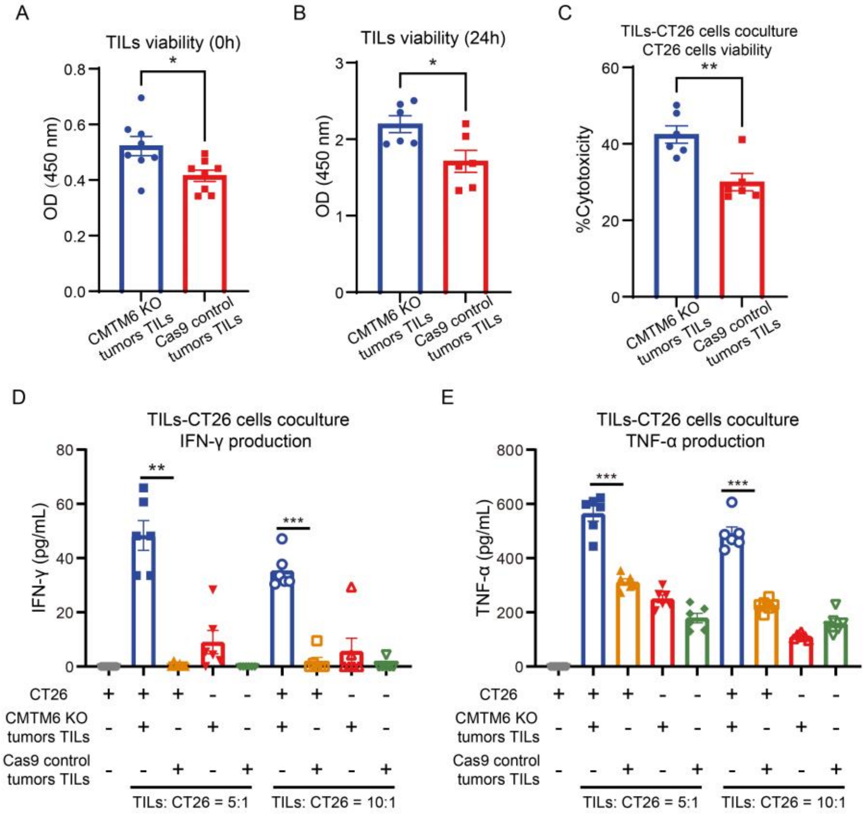


**Fig. S6. Depletion of CMTM6 enhanced specific antitumor cytotoxicity.** At 12 days after tumor bearing, TILs in CMTM6-deficient tumors and control tumors were isolated. (**A** and **B**) TILs were detected for cell viability by CCK8 at 0 h (A; n = 8) and 24 h (B; n = 6). (**C** to **E**) CT26 cells and TILs were co-cultured at 5:1 and 10:1 ratio for 24 h, respectively (n = 6). The viability of CT26 cells (5:1; C) and the concentration of IFN-γ (D) and TNF-α (E) in cell supernatant were determined. The data are presented as the mean ± SEM. * $p < 0.05$; ** $p < 0.01$; *** $p < 0.001$ by unpaired t test or one-way ANOVA followed by Tukey's multiple comparisons test.

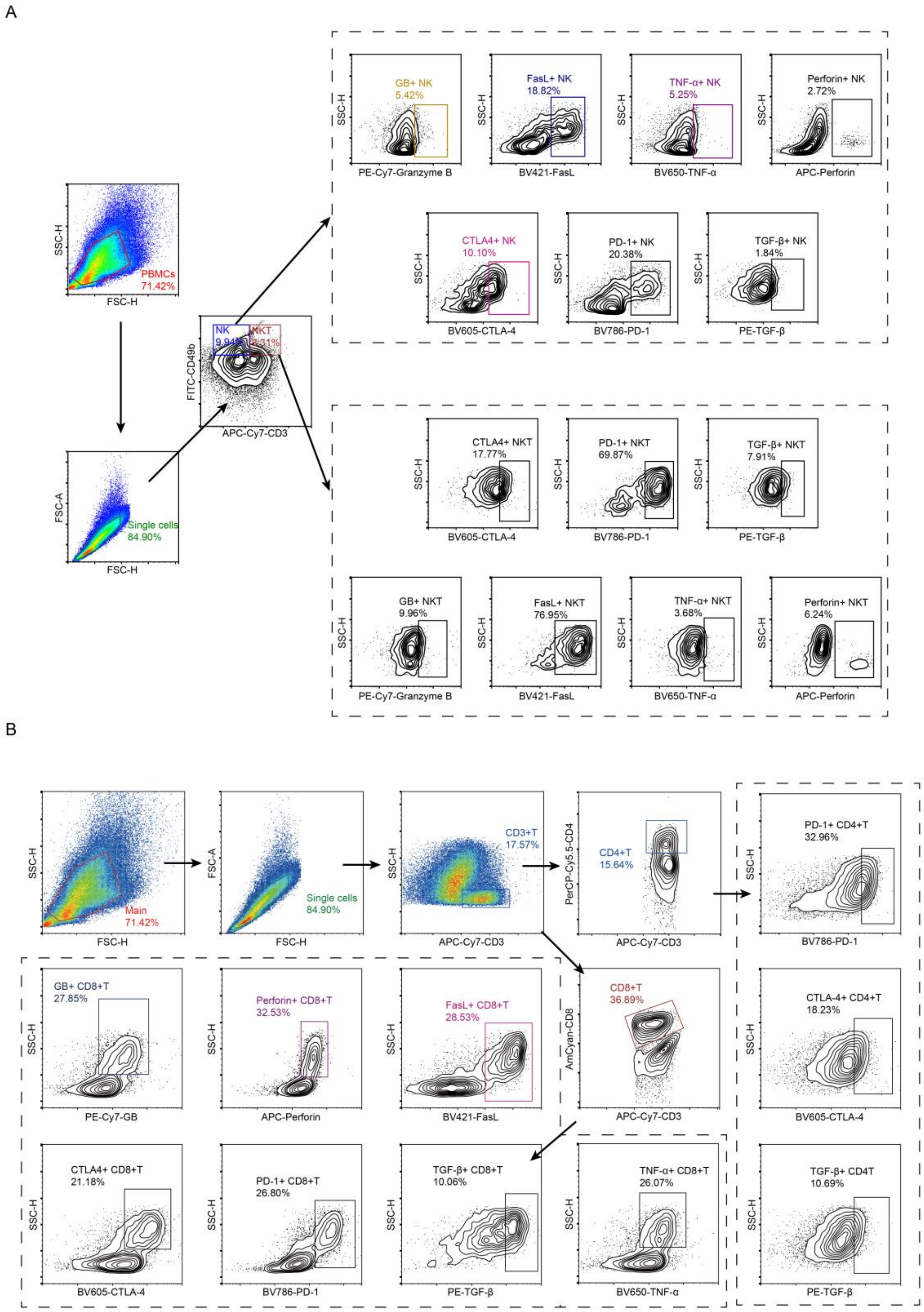


**Fig. S7. Flow-cytometry gating strategy for quantifying lymphocyte subsets in murine tumors.** (**A**) Representative flow-cytometry gating strategy for quantifying the NK and NKT cell subsets in murine tumors. (**B**) Representative flow-cytometry gating strategy for quantifying the $CD4^+$ T-cell and $CD8^+$ T-cell subsets in murine tumors.

A

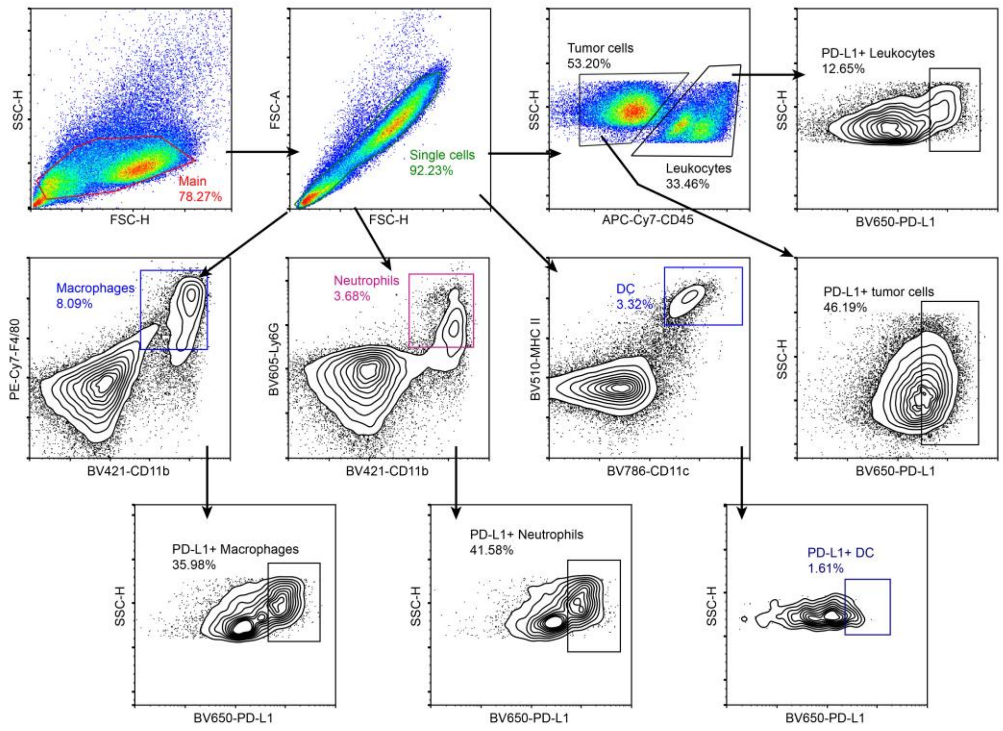


B

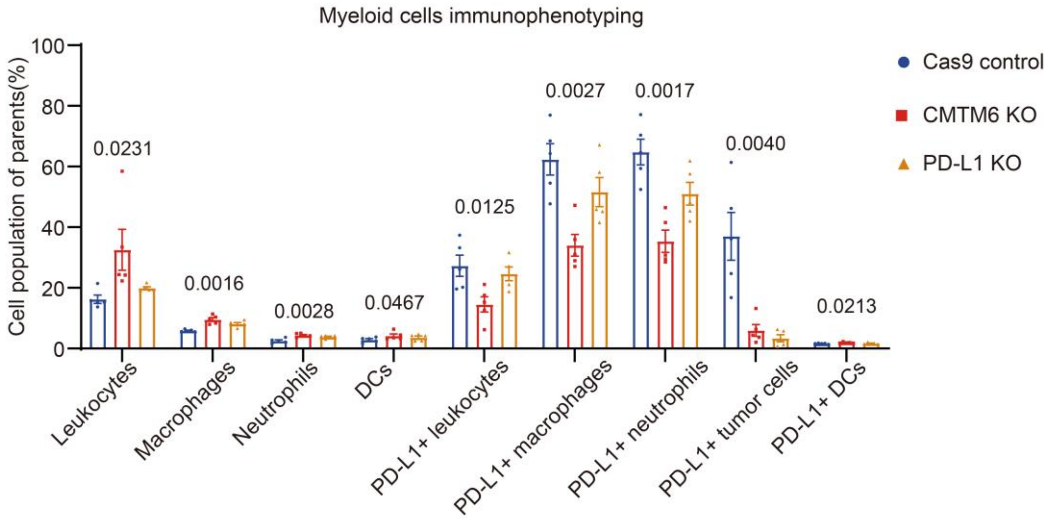


**Fig. S8. Flow-cytometry gating strategy for quantifying myeloid cells in murine tumors.** (**A**) Representative flow-cytometry gating strategy for quantifying the non-lymphocyte subsets in murine tumors. (**B**) The histogram shows the immunotyping of intratumoral infiltrated myeloid cells from CMTM6-deficient CT26 tumors, PD-L1 deficient CT26 tumors and control CT26 tumors by flow cytometry (n = 5). The data are presented as the mean ± SEM and are analyzed by one-way ANOVA followed by Tukey’s multiple comparisons test.

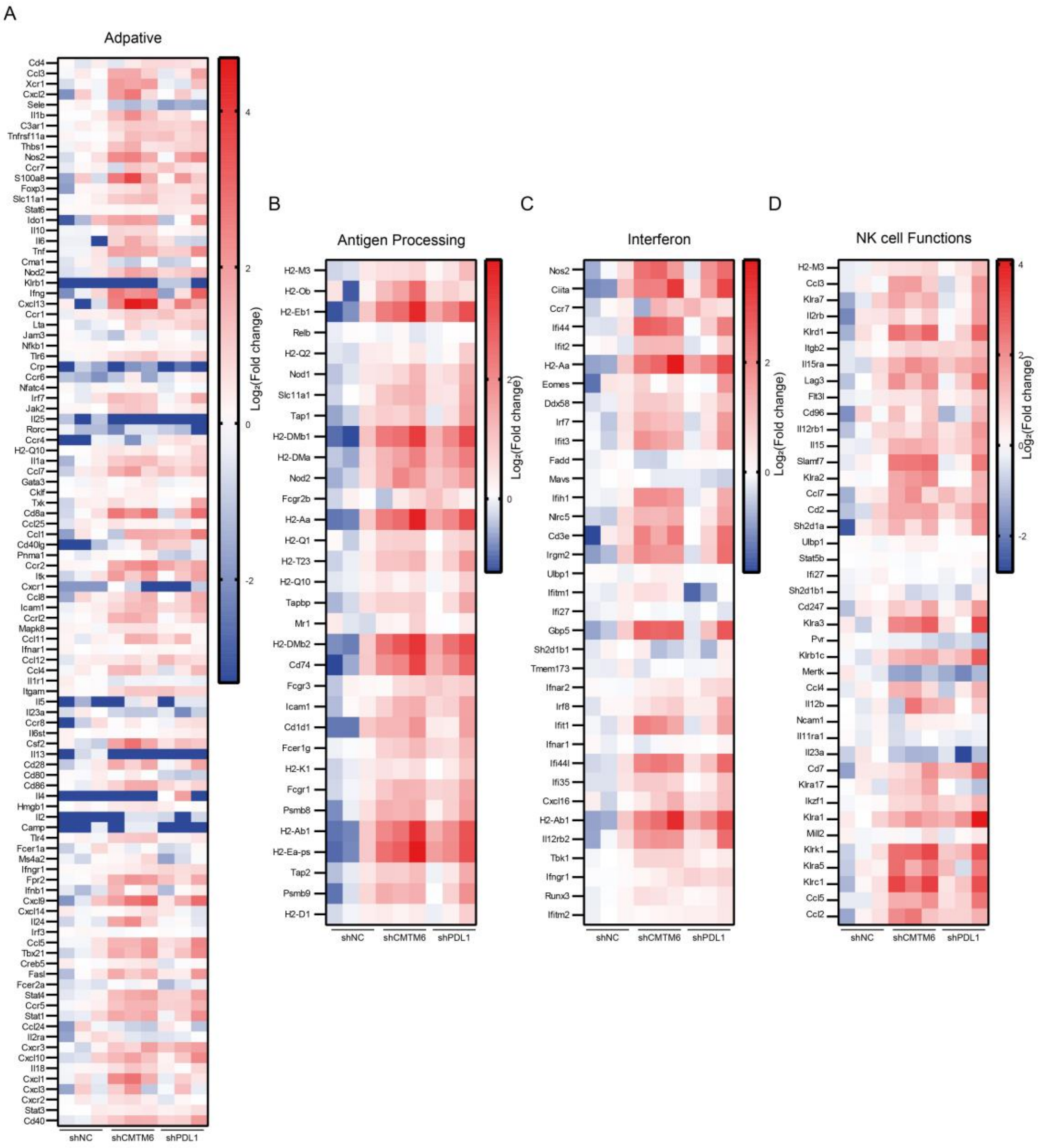


**Fig. S9. Tumor CMTM6 knockout affected intratumoral adaptive immune responses, antigen processing, interferon process and NK cell functions.** (**A** to **D**) RNA-sequencing analysis of tumor tissues from BALB/c mice bearing with shNC, shCMTM6 or shPD-L1 CT26 tumors (n = 3). Heatmaps show that immune pathway analysis (adaptive immune responses, antigen processing, interferon process and NK cell functions). Gene expression (FPKM) was calculated as the $\log_2$(fold change) compared to the shNC group.

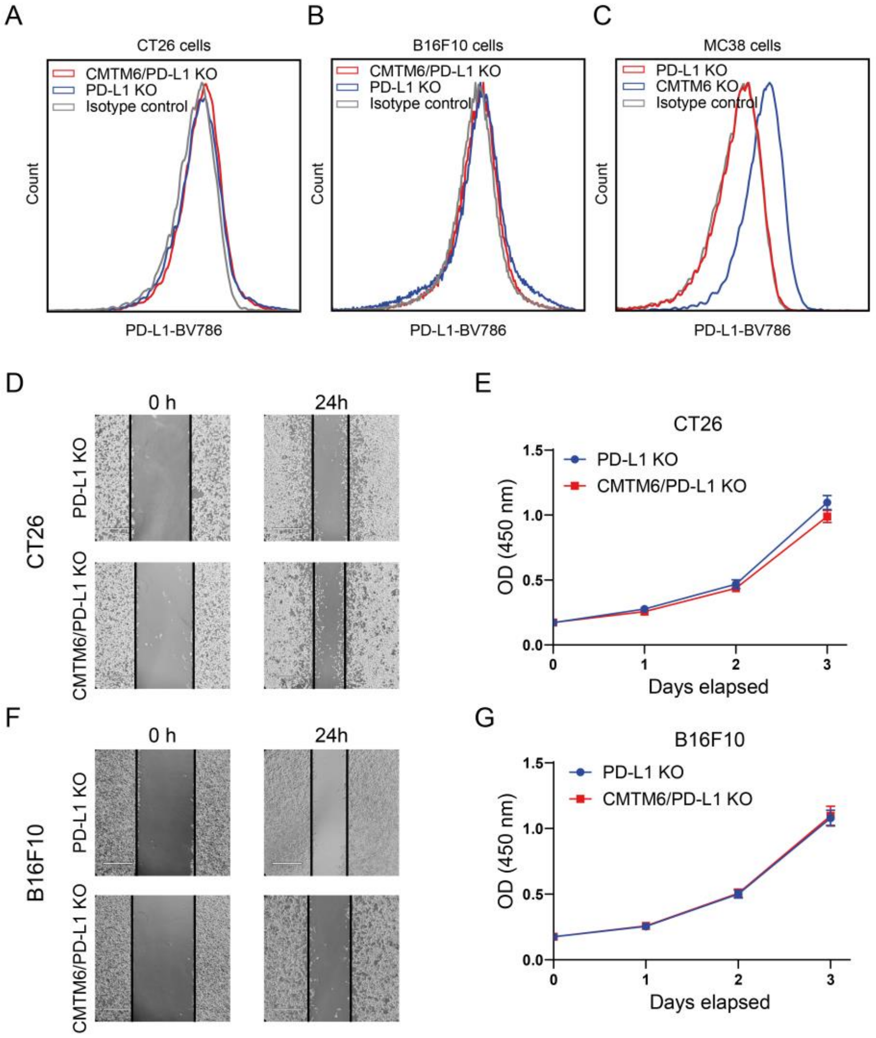


**Fig. S10. Knockout of CMTM6 and PD-L1 and its effect on tumor-cell growth in vitro.** (**A** to **C**) Flow cytometry analysis of the expression of PD-L1 in CMTM6-deficient murine tumor cells with PD-L1 knockout. (**D** to **G**) The wound-healing assays (n = 3) and cell proliferation assays (n = 6) were performed on murine tumor lines with CMTM6 and PD-L1 knockout. The data are presented as the mean ± SEM.

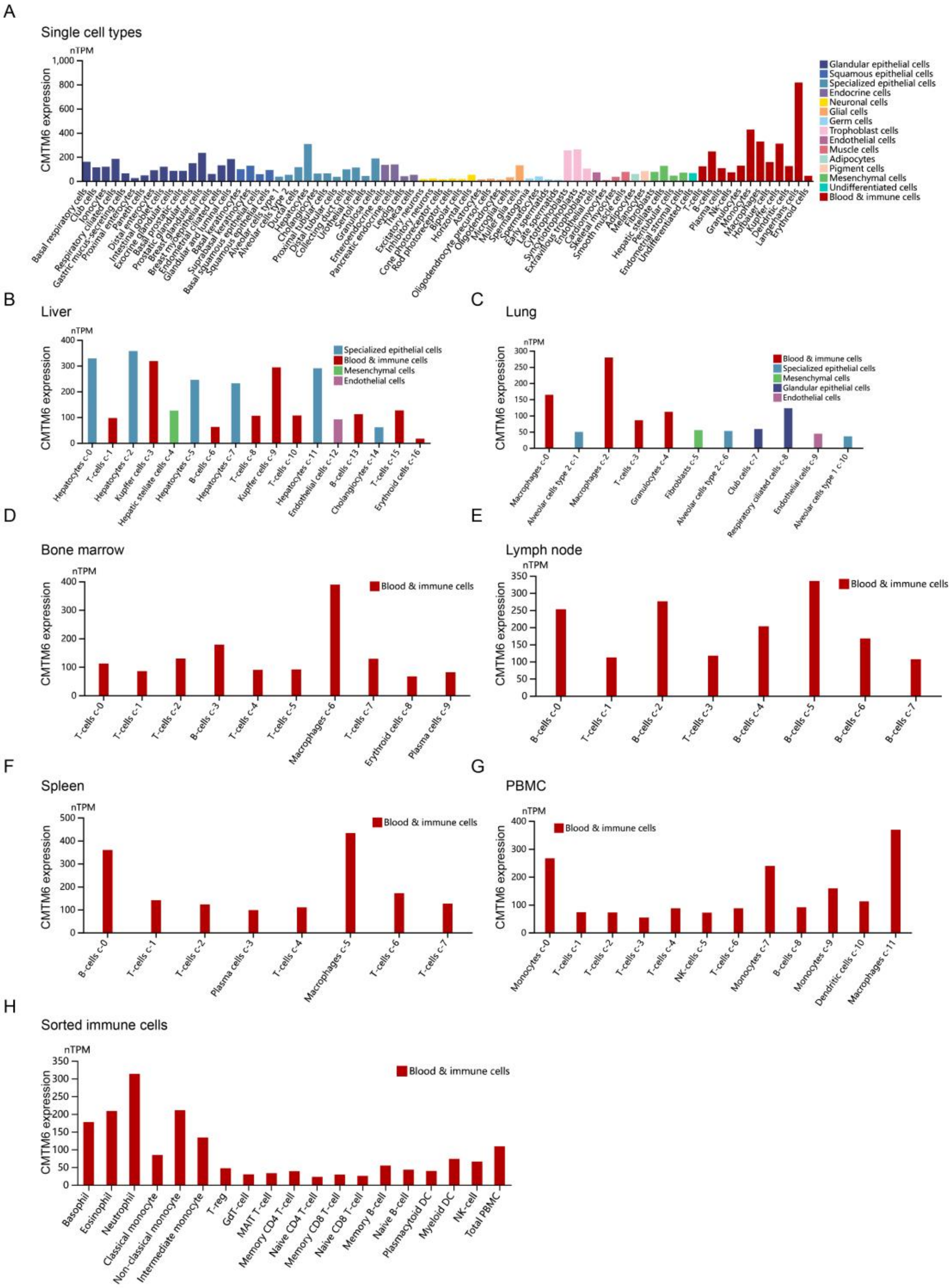


**Fig. S11. Expression profile of CMTM6 at the single cell level. (A** to **H)** At the single-cell transcription level, the expression of human CMTM6 in tissues of the whole body, as well as in liver, lung, bone marrow, lymph node, spleen and PBMC was analyzed by the Human Protein Atlas Project (https://www.proteinatlas.org/).

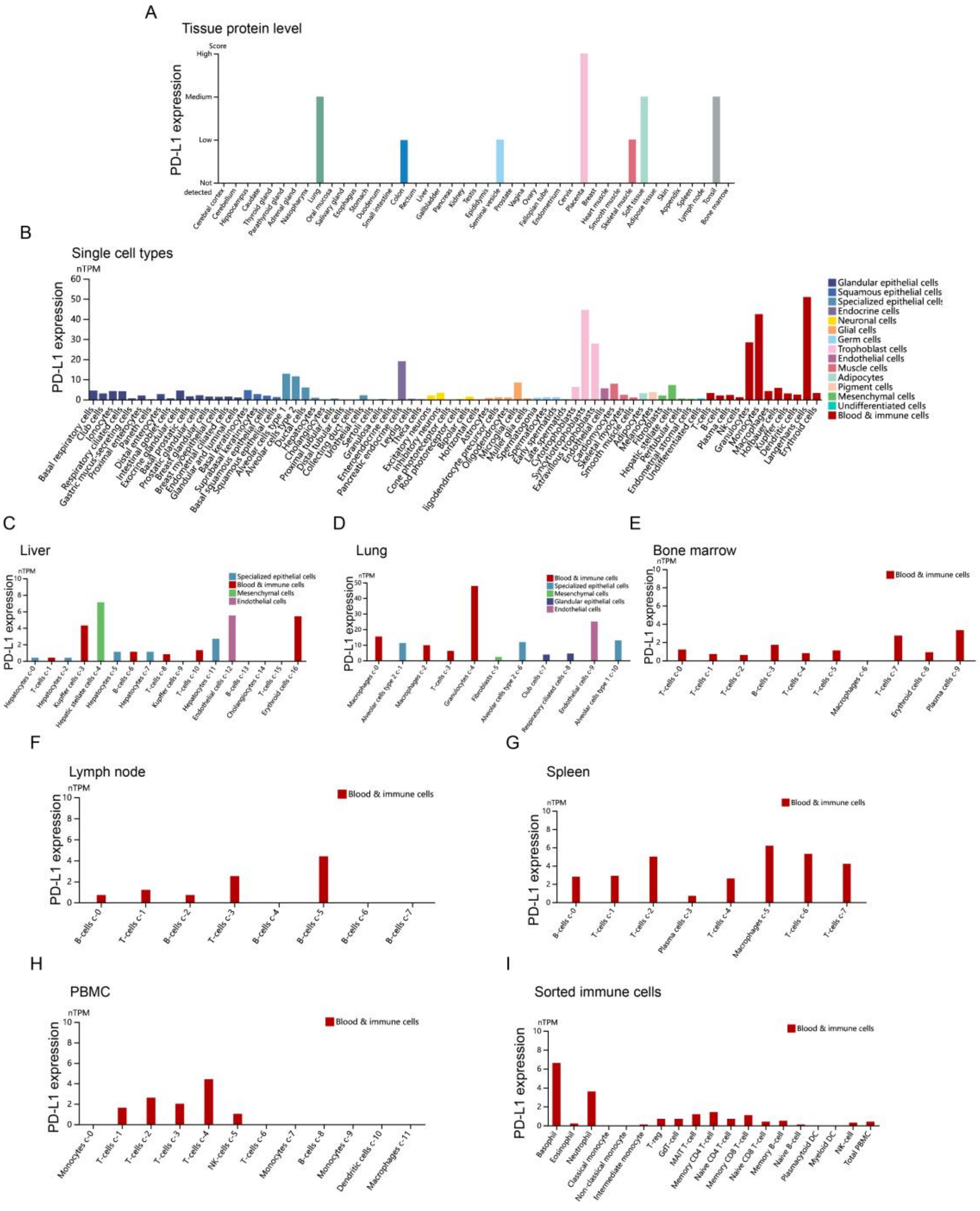


**Fig. S12. Expression profile of PD-L1 at the single cell level. (A)** At the protein level, the expression of human PD-L1 in tissues of the whole body was analyzed by the Human Protein Atlas Project (https://www.proteinatlas.org/). **(B** to **I)** At the single-cell transcription level, the expression of human PD-L1 in tissues of the whole body, as well as in liver, lung, bone marrow, lymph node, spleen and PBMC was analyzed by the Human Protein Atlas Project.

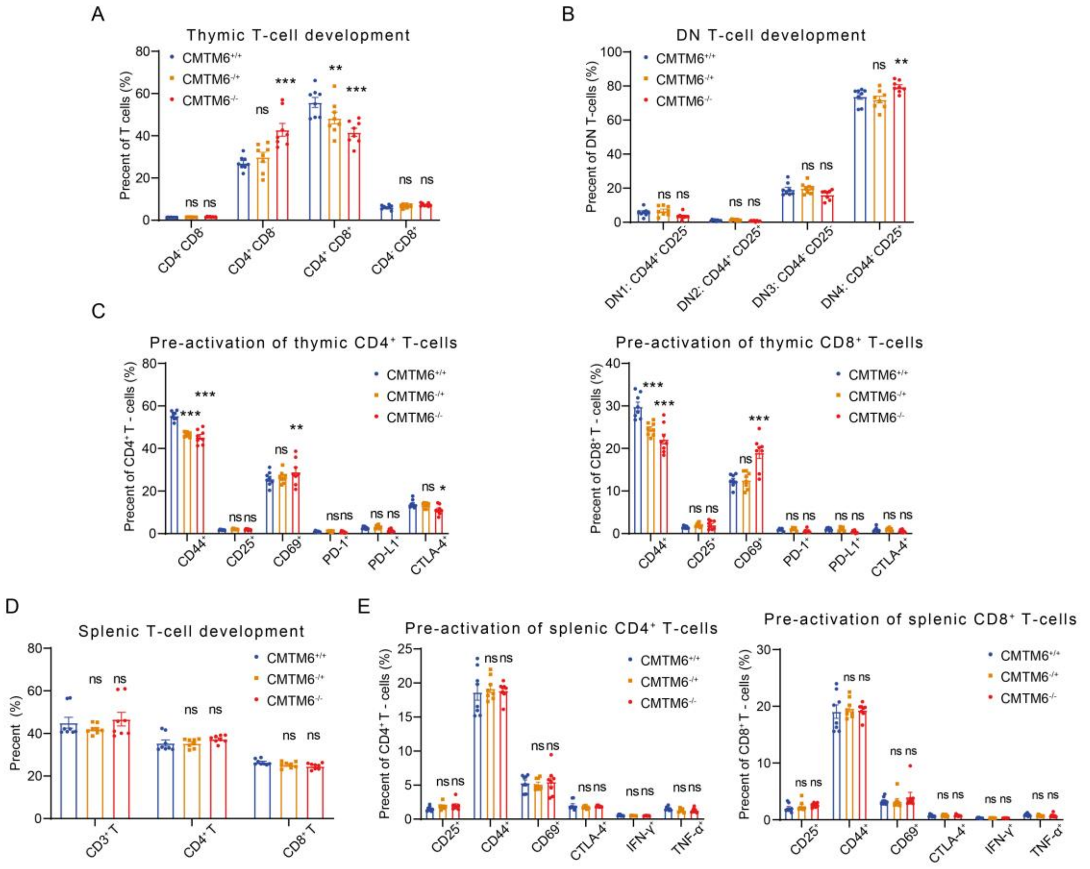


**Fig. S13. CMTM6 affected T cell development and pre-activation.** Without stimulation, splenocytes and thymocytes were detected for pre-activated phenotypes and cell development by flow cytometry. (**A**) Thymic $CD3^+$ T-cells from $CMTM6^{-/-}$, $CMTM6^{-/+}$ and $CMTM6^{+/+}$ mice were tested for expression of CD3 and CD8 (n = 8). (**B**) Thymic $CD4^–$ $CD8^–$ T-cells from $CMTM6^{-/-}$, $CMTM6^{-/+}$ and $CMTM6^{+/+}$ mice were tested for co-expression of CD44 and CD25 (n = 8). (**C**) $CD4^+$ and $CD8^+$ thymic T-cells from $CMTM6^{-/-}$, $CMTM6^{-/+}$ and $CMTM6^{+/+}$ mice were tested for expression of activation markers and checkpoints (n = 8). (**D**) The proportion of T-cells in the spleen from $CMTM6^{-/-}$, $CMTM6^{-/+}$ and $CMTM6^{+/+}$ mice was analyzed (n = 8). (**E**) $CD4^+$ and $CD8^+$ splenic T-cells from $CMTM6^{-/-}$, $CMTM6^{-/+}$ and $CMTM6^{+/+}$ mice were tested for expression of activation markers, checkpoints and cytokines (n = 8). The data are presented as the mean ± SEM. * $p < 0.05$; ** $p < 0.01$; *** $p < 0.001$; ns not significant by two-way ANOVA followed by Tukey’s multiple comparisons test.

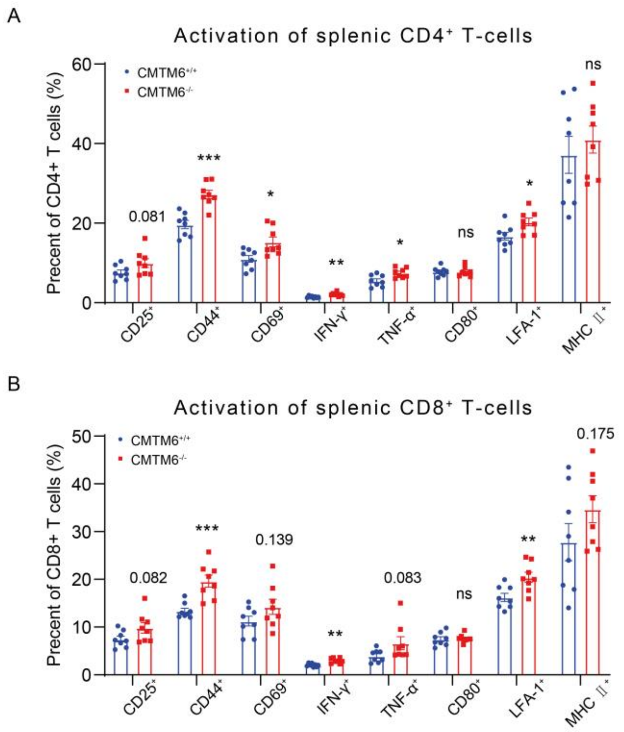


**Fig. S14. CMTM6 affected T cell activation.** Splenocytes from CMTM6$^{-/-}$ and CMTM6$^{+/+}$ mice were stimulated with PMA and ionomycin for 24 h for activation phenotype analysis. (**A**) CD4$^{+}$ splenic T-cells were tested for the expression of surface activation markers and cytokines (n = 8). (**B**) CD8$^{+}$ splenic T-cells were tested for the expression of surface activation markers and cytokines (n = 8). The data are presented as the mean ± SEM. * $p < 0.05$; ** $p < 0.01$; *** $p < 0.001$; ns not significant by two-way ANOVA followed by Tukey's multiple comparisons test.

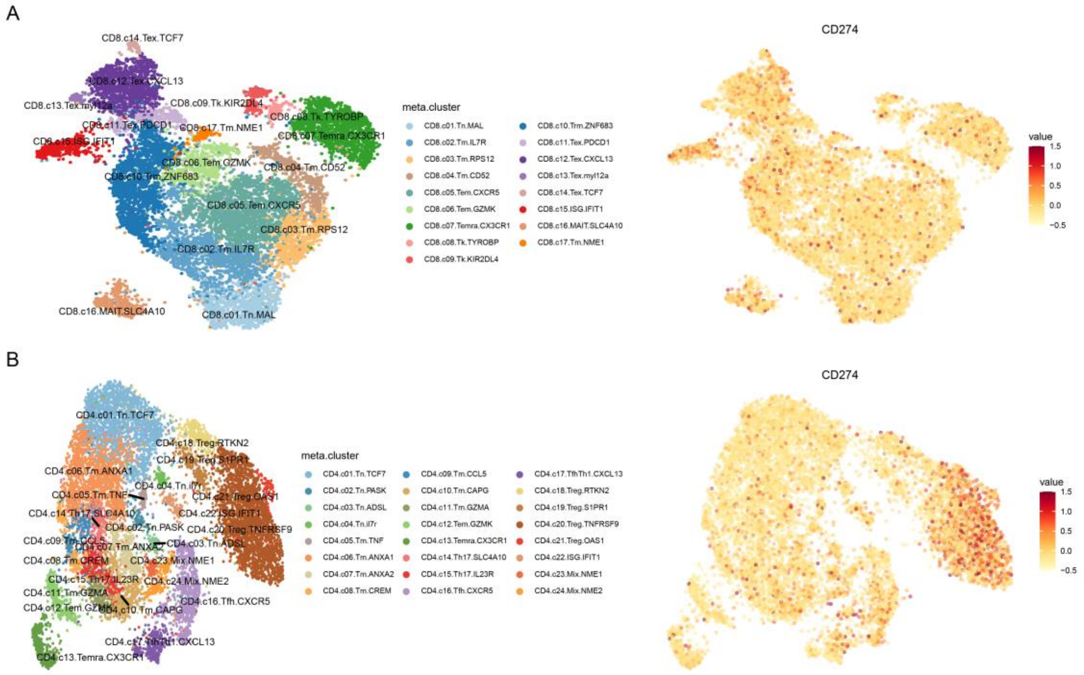


**Fig. S15. Pan-cancer T-cell expression profile of PD-L1.** (**A** and **B**) Single cell RNA-seq data for PD-L1 expression in tumor-infiltrating CD8$^+$ (A) and CD4$^+$ (B) T-cells were obtained from the GEO database (GSE156728) and analyzed by the scDVA interactive tool (http://cancer-pku.cn:3838/PanC_T/).

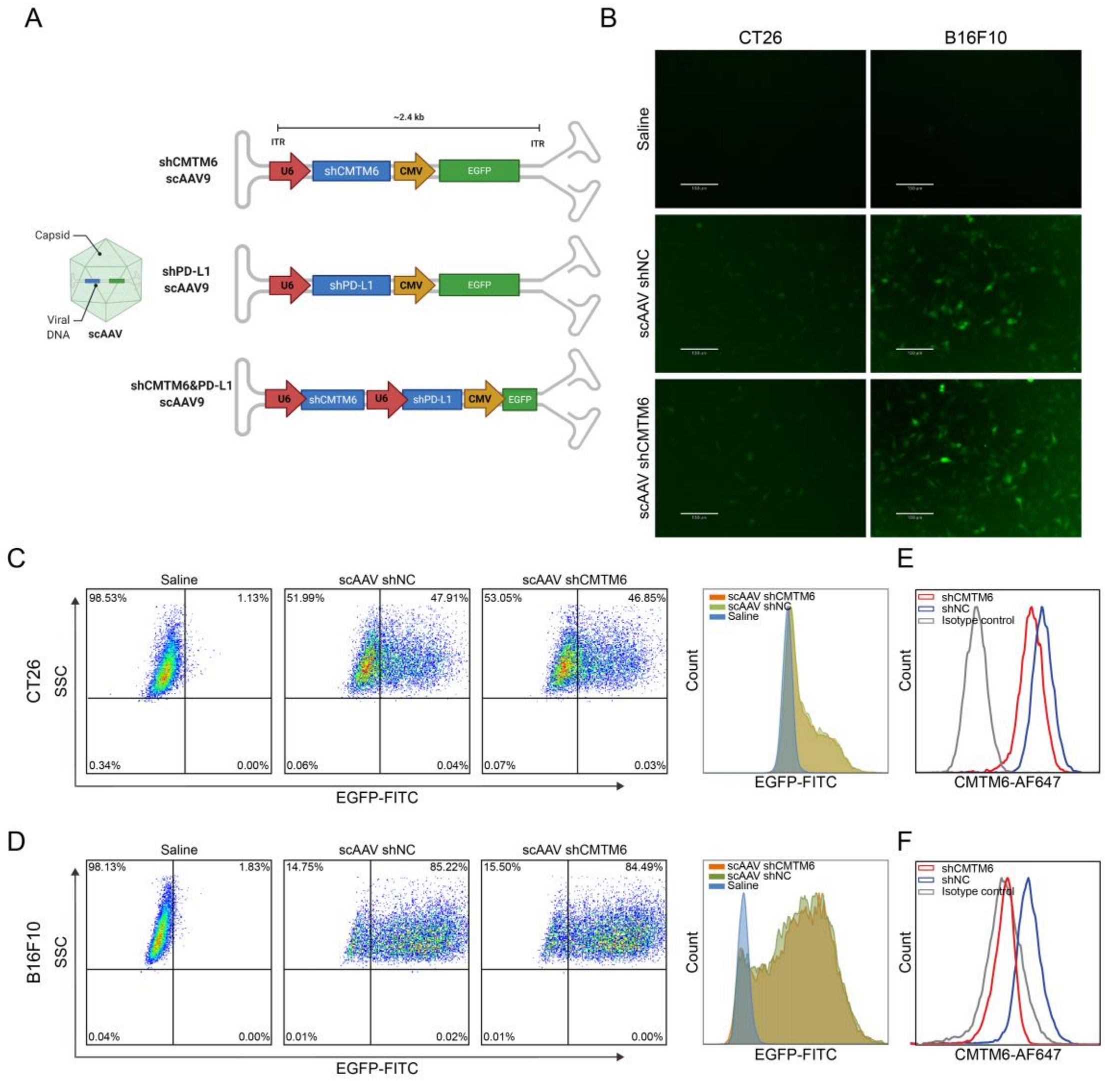


**Fig. S16. scAAV effectively infected mouse tumor cells in vitro.** (**A**) Illustration of scAAV vectors used to deliver the shRNA expression in this paper. (**B** to **F**) scAAV infected CT26 cells and B16F10 cells with a MOI of 10,000. After 72 hours of infection, the green fluorescent signal was detected by flow cytometry (n = 3) and fluorescence microscopy (n = 3). The CMTM6 expression was detected by flow cytometry (E and F; n = 3).

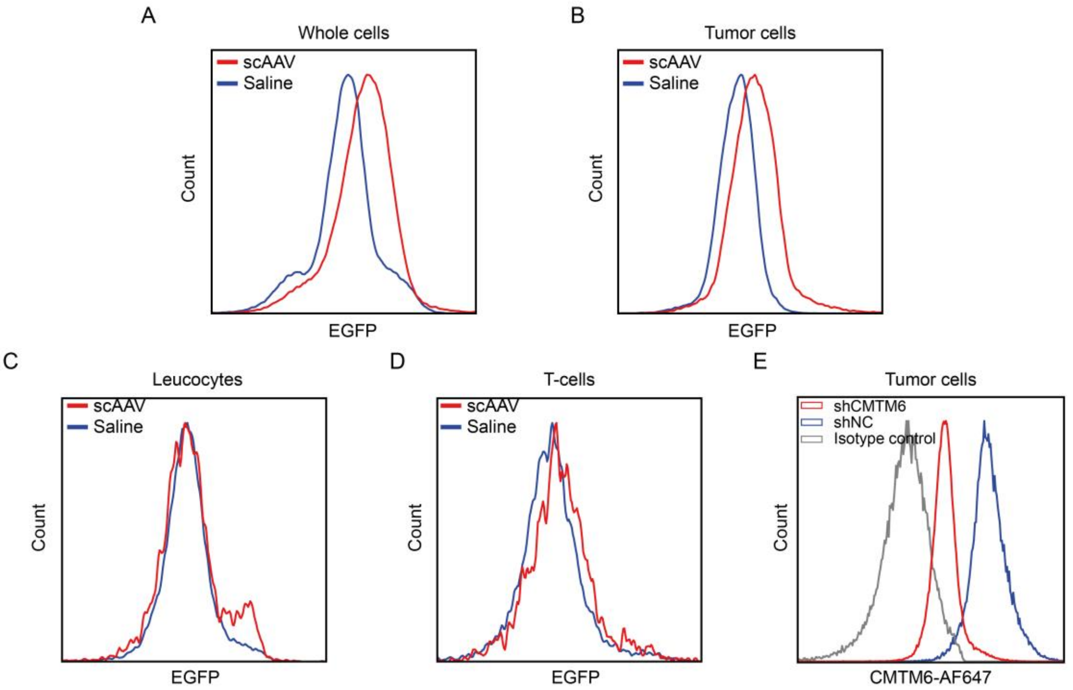


**Fig. S17. scAAV effectively infected mouse tumor cells in vivo.** (**A** to **E**) $5\times10^9$ of scAAV-shCMTM6 was peritumorally injected on day 6 of CT26 tumor growth (n = 3). After 3 days, the tumor tissues were collected and processed into single cell suspensions, and the green fluorescence signal in each cell subpopulation was analyzed by immunotyping. And the CMTM6 expression on tumor cells was detected by flow cytometry (E).
